# Five-layered robot (F-robot) based on functionally graded multilayer foam antiviral low-carbon (F-MAX) system for Pathogenic Microorganism Elimination

**DOI:** 10.1101/2024.04.15.589384

**Authors:** Haotian Fan, Wangcheng Gu, Dongrui Zhou, Song Ge, Pengfeng Xiao, Zhongjie Fei

## Abstract

In this study, a revolutionary air filtration technology, the F-MAX multilayer composite plate, is introduced, offering high efficiency and environmental sustainability. This innovative system is designed to capture a wide range of pollutants, including harmful viruses and bacteria, enhancing air quality significantly. The F-MAX combines multiple layers, each tailored to target specific particles, with features like an electrostatically charged melt-blown fabric and eco-friendly materials like lithium brine by-product magnesia. Its durability, antiviral, and antibacterial properties make it a sustainable choice for air purification, suitable for both commercial and residential use. This system represents healthier living environments, effectively removing airborne contaminants, and demonstrating a commitment to a sustainable future. Additionally, the study introduces the F-robot specifically designed for laboratory environments to ensure pristine air quality.

**One-Sentence Summary:** T-robot air filter, which use F-MAX, a multilayer composite consisted of self-healing cellular coating, Desert Rose (DR) coating, melt-blown cloth, and BMSC with high-efficiency, environmentally sustainable filtration, and antiviral properties, suitable for diverse environments.

---

In the modern interconnected world, diseases can swiftly traverse borders and continents, wreaking havoc on both public health and the global economy. Thus, the threat of disease outbreaks looms larger than ever before. This challenge is compounded by the virulent nature of viral infections and their diverse modes of transmission—whether aerosols, droplets, or everyday surfaces (fomites). The H1N1 pandemic has magnified awareness of the perils posed by high-touch surfaces. Thus, innovative solutions have been explored to curb the presence of active viral pathogens. Another challenge in the modern world is the livestock and poultry farming industry, which is progressively moving toward more intensive and large-scale operations. However, the growth in animal stocking density has also led to an increased risk of disease among farmed animals, making disease prevention and control considerably more challenging. Surveys have indicated that over 70% of animal diseases are predominantly viral in nature (*1*). Viral disease outbreaks in animals will adversely impact the quality of livestock and poultry products, and the livestock industry will experience significant animal losses and substantial economic setbacks. Moreover, some zoonotic viruses, which are viruses that originate from farm animals, have the potential to cross over into human populations, posing a direct threat to human health. For instance, the global H1N1 influenza A viral outbreak in 2009 (*2*) and the spread of the highly pathogenic avian influenza H5N2 subtype in the United States in 2014–2015(*3, 4*) were both large-scale epidemics caused by zoonotic viruses originating from farm animals. Therefore, controlling viral contamination and outbreaks in livestock and poultry farms is crucial for combatting the spread of disease. The use of appropriate facility design to implement preventative measures at the source is a fundamental strategy for managing the transmission of viruses to humans and mitigating the initial outbreak of epidemics.

Microbes can survive on a variety of different surfaces, both in healthcare settings and on common surfaces. This is often a crucial vector of transmission for infectious diseases. Therefore, strategies for preventing bacterial transmission and biofilm formation by killing and/or reducing the attachment of microbes to surfaces have been widely researched. For example, the use of biocidal coatings, surface-bound active antimicrobials, or passive pathogen-repellent surfaces have been designed using chemical modification, nanomaterials, and micro- and nanostructuring.

Air pollution contains various harmful chemical compounds. These compounds include small, solid particles known as particulate matter, which can be classified as PM2.5 (2.5 µm) or PM10 (10 µm) (*5*). Viruses (*6*), bacteria, fungi, and inorganic substances such as metal ions, salt, and other solids (*7*) can all be harbored by airborne PM2.5 particles. Moreover, the small size of PM2.5 particles means they cannot be easily removed through natural inhalation (*8*). Consequently, these particles frequently penetrate the respiratory system (*9*), where they pose a particular hazard (*10*). Specifically, PM2.5 particles are easily inhaled directly into the lungs, leading to health issues such as alveolar tissue damage (*11*), eye and nasal irritation, respiratory tract infections, and vision impairment (*12*). Consequently, air filtration is extremely important for enhancing the quality of inhaled air (*13*).

Poor outdoor air quality also negatively impacts indoor air quality (*14*). The quality of air is defined as poor if the concentration of pollutants, including solids and gases, surpasses the healthy air threshold (*15*). Gas-phase compounds such as CO, NO, SO, and HC can disrupt the human respiratory system, leading to mucosal irritation and heart problems. Moreover, these gases significantly contribute to the global greenhouse effect (*16*).

Many common materials have traditionally been employed in air purification and filtration applications, including paper, cotton, activated carbon, and fiberglass. These materials exhibit distinct advantages as well as drawbacks. For instance, benefits such as cost-effectiveness, facile scalability in production and replacement, ready accessibility, consistent high-quality output, lightweight properties, ease of manipulation and replacement, and wide-ranging applicability can be achieved using traditional air filtration materials. However, these materials also have inherent limitations in filtration efficacy, such as their inadequate performance for the removal of tiny particles and noxious gases, inability to isolate viruses and bacteria, and suboptimal air permeability efficiency. Moreover, these materials need to be frequently replaced. Some traditional filtration materials might require chemical additives or emit harmful substances during manufacturing, leading to potential environmental hazards. Overall, traditional air filtration materials represent an economical and readily accessible option for fundamental air filtration requirements. However, these traditional materials are not suitable for applications with more stringent air quality mandates, such as the elimination of fine particulate matter and specific hazardous gases. To meet these requirements, advanced filtration technologies are imperative. Thus, highly efficient and multifunctional filtration materials with low carbon footprints are increasingly in demand.

Air filter media typically consist of synthetic fiber polymers prepared using petroleum by-products. Polypropylene, polyester, polyamide, and glass fibers are some of the commonly used synthetic polymer fibers (*17*). However, these synthetic fiber polymers exhibit several drawbacks compared to natural polymer fibers derived from plant sources (*18*). For instance, petroleum-derived polymer fibers are prepared using complex production processes, and they cannot be biodegraded or recycled (*19*). Additionally, the production of these fibers leads to greenhouse gas emissions (*20*). In contrast, natural fibers such as cellulose (*21*) offer numerous advantages. Because they grow naturally, natural fibers are more accessible, and they can be prepared using cost-effective production methods (*22*). Moreover, natural fibers are environmentally friendly (*23*), and they can be used to absorb air pollutants (*24*).

The onset of the pandemic (*25*) has led to significantly increased demand for air filter media (*26*), particularly for applications such as face masks(27). Thus, raw materials that are safe, inexpensive, and easily processed are required (28). However, very few investigations have investigated the development of materials with antiviral, filtration, and building material functions. In this study, a new kind of multilayer plate(F-MAX) with a low gravity, price, and carbon emissions was developed, and being used in the newly developed F-robot which is an active air purifier.

This plate exhibits significant anti-virus performance and can be used in air filtration applications. This new technology successfully balances the trade-off between antiviral performance, low carbon, and excellent comprehensive physical characteristics. As shown in Fig. 1, this symmetric plate is composed of five layers in the following order:, a glass fiber (GF) net with magnesium phosphate cement (MPC) foaming paste, an electrostatically charged melt-blown fibrous filter, another GF net glued on another FBMSCP plate painted with, final layer, cellular coating With extreme high resistance against environmental effects.

**Figure 1.**
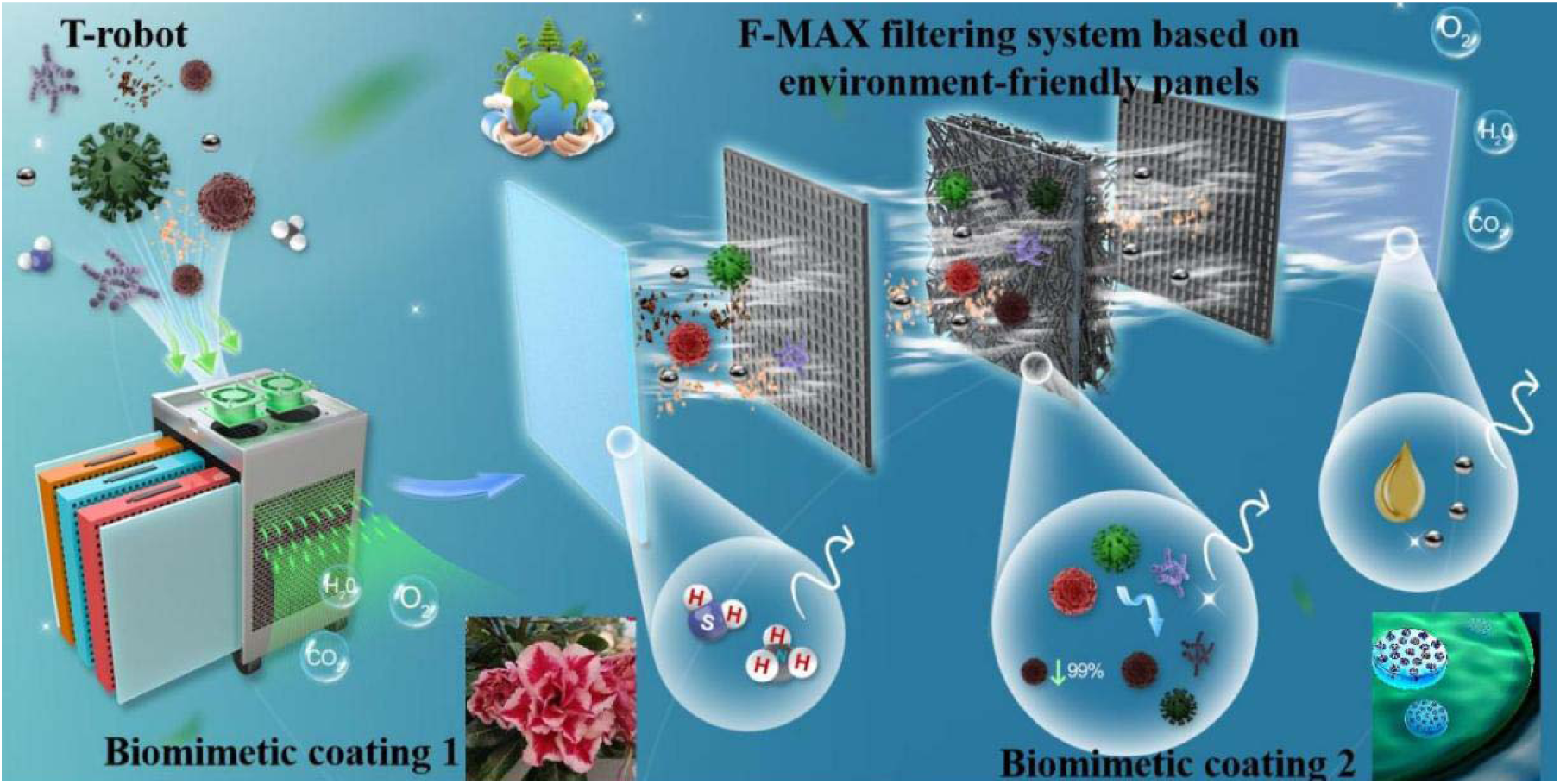
The depicts the overall structure of the F-robot, which is based on the F-MAX filtering system. The F-MAX plate demonstrates significant antiviral performance and is suitable for air filtration applications. This new technology effectively balances the trade-off between antiviral performance, low carbon emissions, and excellent overall physical properties. The symmetric plate shown in the figure consists of five layers: magnesium phosphate cement (MPC) foam paste with glass fiber (GF) net, covered with DR coating, a newly developed superhydrophobic material, an electrostatically charged melt-blown fibrous filter, another GF net adhered to another FBMSCP plate, and a final layer of cellular coating, which provides extremely high resistance to environmental effects.

Comprising multiple specialized layers, F-MAX combines a foaming basic magnesium sulfate cement paste (FBMSCP) and a hydrophobic Desert Rose coating for robust particle filtration. Its centerpiece is an electrostatically charged melt-blown fabric, crucial for trapping fine particles and microorganisms, and surpassing the standards set for surgical masks in terms of static charge retention and antiviral capabilities.

F-MAX’s sustainable design is one of its key advantages, featuring eco-friendly materials like lithium brine by-product magnesia and flue gas desulfurization by-product sulfuric acid, contributing to a low carbon footprint and high dust holding capacity. The filter’s durability is enhanced by ultra-durable superhydrophobic cellular coatings, ensuring long-term performance even under demanding conditions.

Capable of filtering various pollutants and effective across a range of particulate sizes, F-MAX is versatile, suitable for both commercial and residential environments. In essence, F-MAX’s innovative design, efficiency, and sustainability mark it as a pivotal solution in air quality management and a significant player in the future of air purification.

## Results

### Filtration efficiency of F-MAX

The actual filtration efficiency of the prepared composite multilayer plate is shown in the following Table 1. As can be seen, the requirements of European standard EN-779 F8-grade filters are met.

**Table 1.**
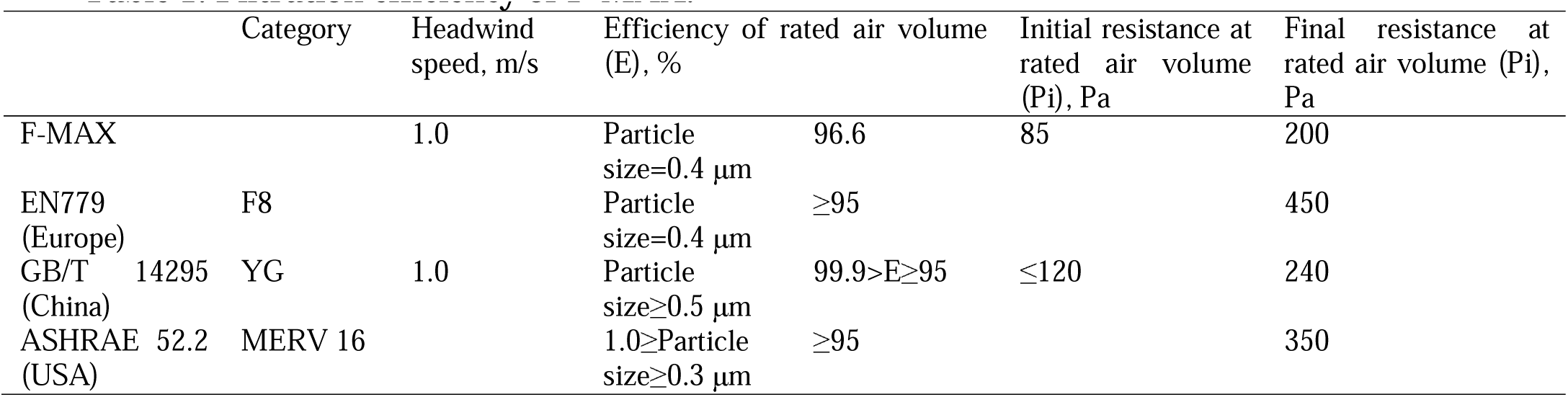
Filtration efficiency of F-MAX.

The effective filtration of impurities in air is realized via the hierarchical combination of a variety of biomimetic materials. As shown in Fig. 2(a), in standard filter media filtration performance experiments, the filtration efficiency of each monolayer material decreases with increasing filter wind speed. Meanwhile, their wind resistance rapidly increases with increasing filter wind speed. The F-MAX filter cartridge demonstrates exceptional air purification proficiency, as demonstrated by its dual capability to filter particulates and efficiently eliminate hazardous gases. The effects of air-flow velocity on the removal efficiency and pressure drop are investigated.

**Figure 2.**
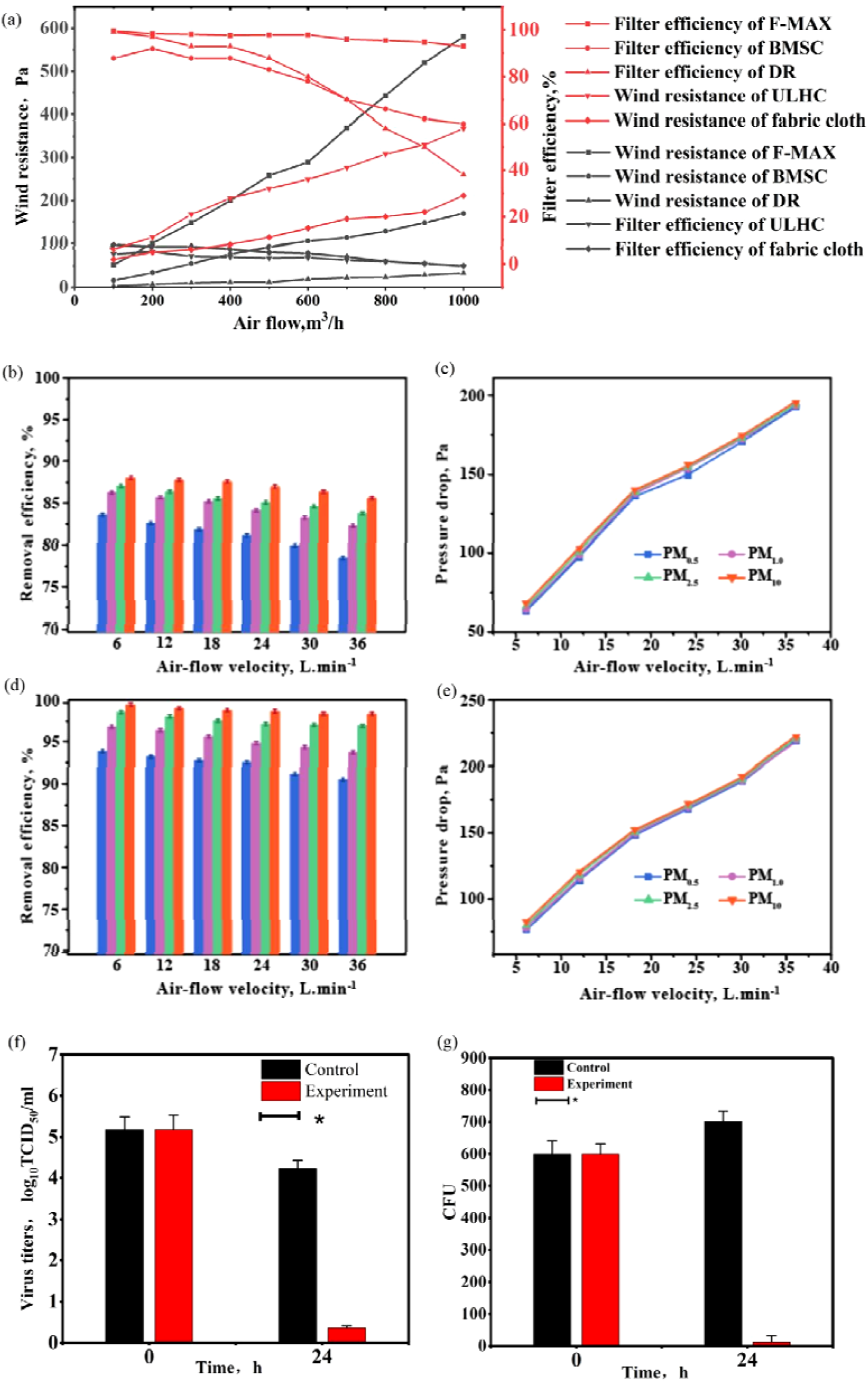
Showcases the superior performance of the F-MAX filter cartridge in air purification. (a) Filtration efficiency and wind resistance, (b–c) Removal efficiency and pressure drop of ordinary commercial filter cartridge. (d–e) Removal efficiency and pressuredrop of F-MAX filter cartridge. (blue, purple, green and orange correspond to PM0.18, PM0.3, PM2.5, and PM10), respectively. (f-g) Antiviral performance of F-MAX filter cartridge. TCID_50_ means 50% tissue culture infective does.

As illustrated in Fig. 2(b), the removal efficiencies of PM based on ordinary commercial filter cartridge of every size are less than 88%. This is mainly because the pores formed by it are too large to block small particles. But they endow it characteristics of low pressure drop as well (Fig. 2(c)). While the removal efficiencies of PM based on F-MAX filter cartridge shows evident higher removal efficiency (Fig. 2(d)), but also a higher pressure drop (Fig. 2(e)). The removal efficiencies of PM based onF-MAX filter cartridge illustrates the superior performance. A significant reduction in the concentrations of the removal efficiencies of PM based onF-MAX filter cartridge can be observed within 36 min. The initial phase of the filter is characterized by a notable decrease in the levels of PM, indicating its robust and rapid absorption properties. This is an essential feature for scenarios requiring durable filtration solutions. The error bars demonstrate the reliability of this filtration process, with minimal fluctuations observed. Therefore, the cartridge has good long-term consistency. Moreover, no rapid saturation phenomenon can be observed. These results demonstrate that F-MAX is an ideal candidate for environments consistently exposed to noxious gases.

Due to the reasonable combination of different layer materials, F-MAX shows comprehensively improved filtration efficiency as well as good filtration performance under higher filter air speeds. As shown in Fig. 3(a)–(d), the filtration effect of this multilayer composite plate material is achieved via the combination of several principles:

1. Physical filtration principle (including mechanical filtration and microfiltration): The multiple layers of the composite plate have different pore structures and particle size distributions, allowing solid particles of various sizes to be retained. As shown in Fig. 1, mechanical filtration is primarily accomplished through the organic coating material of the first cellular coating surface layer. Moreover, this coating is highly hydrophobic due to the contact angle of it is 125°, which made the coating very good at blocking pathogenic microorganisms. Besides, the coating with “self-healing” design (Fig. 3(a)) which is highly durable has small pores capable of filtering particles with diameters larger than 5 µm, as shown in Fig. 3(a). Thus, the adsorption capacity and dust holding capacity of this layer will not be affected by the adsorption of water in the filtered air. A microfiltration effect is achieved through a combination of the coating, BMSC foamed concrete, and melt-blown fabric. The porous structure of the coating initially adsorbs larger particles. Air carrying smaller particles passes through the coating, where it undergoes a second-stage microfiltration step in the circular pores (size distribution: 1-3 µm) of the BMSC foamed concrete panel. As shown in Fig. 3(d), SEM analysis of BMSC demonstrates the presence of needle-like crystals composed of hydrated magnesium sulfate and magnesium oxide in a 5·1·7 phase. These needles are present in greater numbers around the circular pores, where they significantly enhance the adsorption efficiency of particles larger than 1 µm.
2. After passing through the physical filtration provided by the first two layers of the structure, air passes through the melt-blown fabric, which is a complex fibrous polypropylene mesh (Fig. 3(c)). Most impurities with particle sizes above 0.3 µm will be adsorbed by this fine mesh, achieving further micropore filtration. Finally, the symmetric structure of the multilayer plate provides a secondary physical filtration step with another BMSC foamed concrete prepared with a lower foaming rate. The smaller inner pores of this BMSC layer enable the effective filtration of particles that bypass the preceding filter layers. Prior to the electrostatic polarization of the middle melt-blown fabric layer, this plate can achieve a filtration efficiency of over 80% for particles larger than 0.4 µm, meeting the requirements of an F7-grade filter according to European standard EN779. The single-sided Desert Rose (DR) coating has a large specific surface area due to its multilayer flower-like folded structure (Fig. 3(b)). The fine pores and specific structure of this layer can filter impurities and corrosive ions from the air. This DR coating acts as a redundant filter material for the overall system. Under high air velocities, the filtration system must be capable of handling large air volumes. This single-sided hydrophilic and oleophobic coating is highly suitable as the final filtration layer.
3. Principles of electrostatic adsorption filtration: the melt-blown fabric layer in the middle of the composite plate can be electrostatically polarized to provide an electrostatic adsorption filtration effect, which utilizes charge to adsorb particulate matter. The melt-blown middle layer of the multilayer plate contains metal and other ion particles that can hold a charge for an extended period. The surface static charge distribution of this layer after undergoing static electricity polarization for 14 days is shown in Fig. 4(a–e). This distribution reveals that the static electric voltage on all areas of the surface exceeds 4 kV. The static electric voltage of the central part of the layer even surpasses 9 kV, which significantly exceeds the requirements for static charge levels in the middle melt-blown fabric layer of regular surgical masks according to international standards. In the structural design of this composite material, the magnesium phosphate cement used for fixation around the edges is relatively thin due to its high bending resistance.

**Fig. 3.**
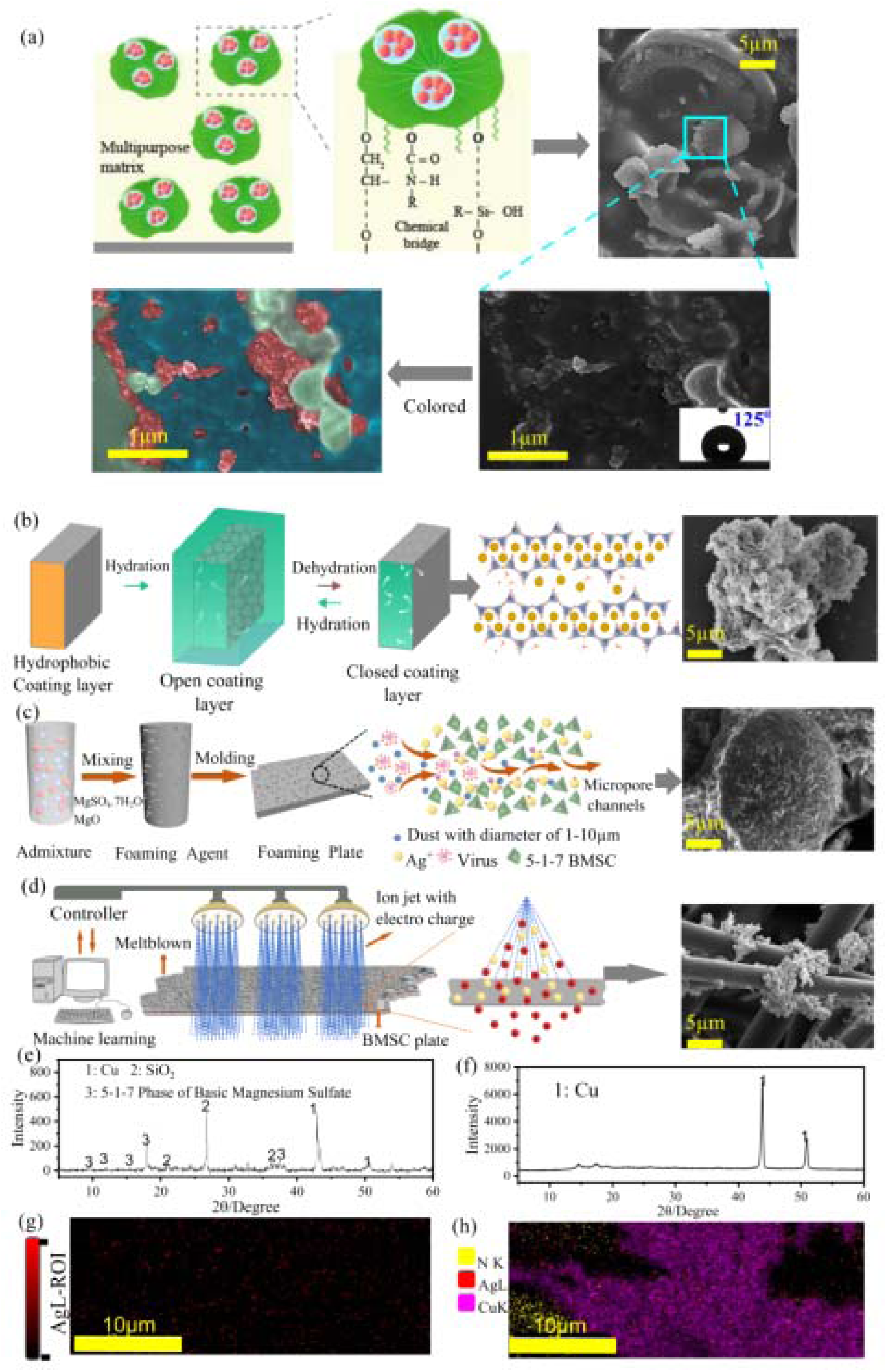
Microscopic analysis of various layered sections in F-MAX. Dssign and SEM image of (a) cellular coating, (b) Desert Rose (DR) coating, (c) melt-blown cloth, and (d) BMSC. (e) XRD patterns of BMSC plate. (f) XRD patterns of melt-blown cloth. (g) EDS mapping of BMSC plate. (h) EDS mapping of melt-blown cloth.

**Fig. 4.**
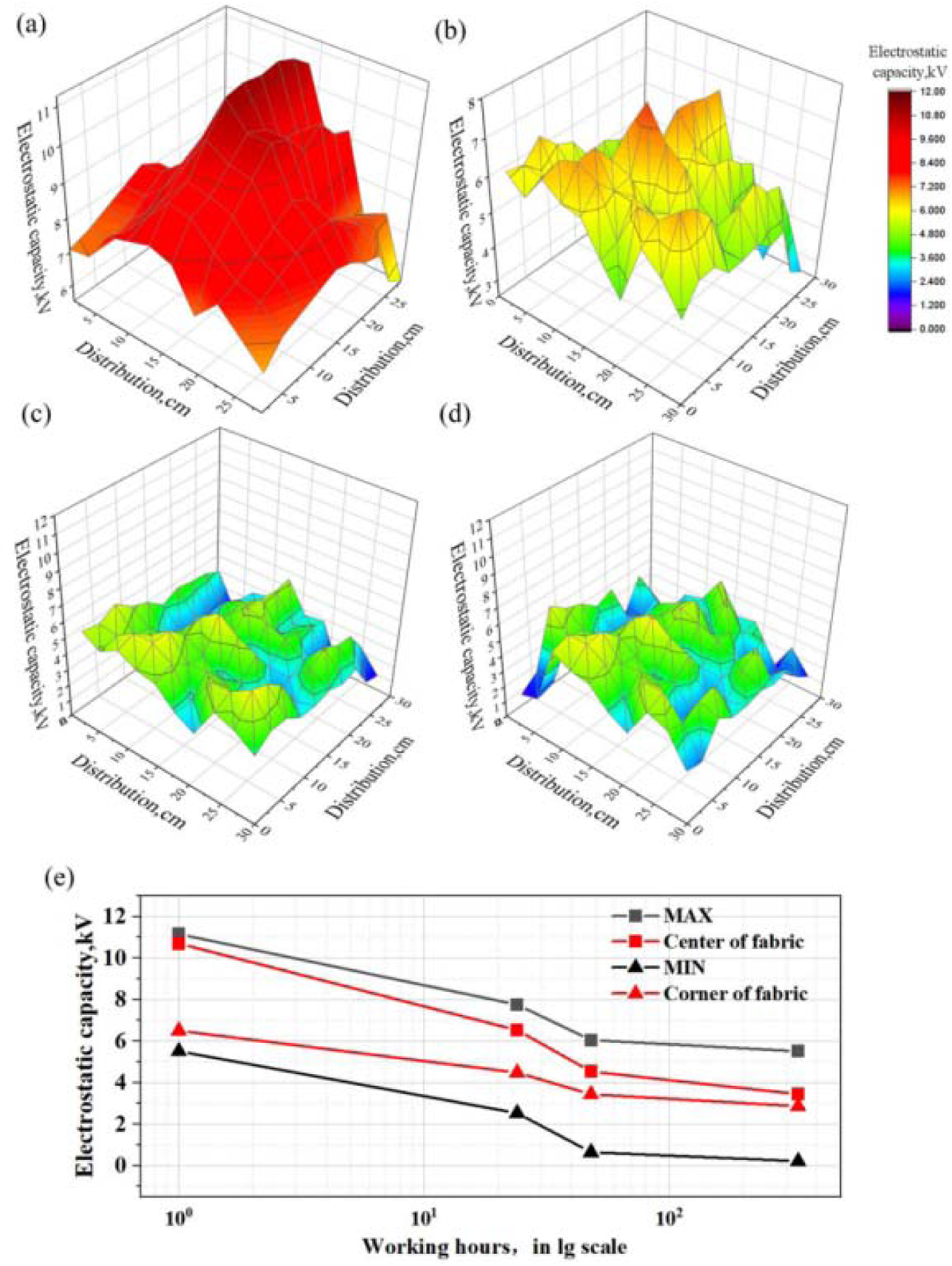
The strong antiviral capabilities of the F-MAX filter over a period of 336 hours (two weeks). It shows that the virus reduction rate starts very high and only slightly diminishes over time, indicating that the F-MAX filter has good long-term reliability and is suitable for extended use.Figure 4(b) displays the antibacterial performance of the F-MAX filter, which like its antiviral capabilities, starts off with a high bacterial reduction efficiency that only slightly declines during long-term operation, affirming its enduring antibacterial efficacy.

Therefore, the multilayer plate can be easily disassembled after the dissipation of static charge in the middle melt-blown fabric layer for re-polarization or replacement of this layer. Experimental results demonstrate that when the middle layer of the multilayer composite plate is electrostatically charged, the achieved filtration efficiency meets the requirements for an F9-grade filter according to European standard EN799.

Figures 4(c) and (d) demonstrate the durability of the F-MAX filter’s cellular coating by showing its resilience when subjected to Taber abrasion cycles. Remarkably, the water contact angle (θ) remains consistently high even after 1000 abrasion cycles, indicating the coating’s strong hydrophobic properties and its ability to maintain performance despite abrasive wear.

### Virus and bacteria elimination efficiency of F-MAX

F-MAX demonstrates outstanding antiviral and antibacterial sterilization and disinfection capabilities. Due to the use of highly wear-resistant hydrophobic coatings on the BMSC plates and DR coatings, bacteria larger than 1 µm are effectively eliminated. Moreover, viruses and bacterial particles are adsorbed on the electrostatic melt-blown cloth. The silver ions and nano-copper particles loaded on the melt-blown cloth then provide antibacterial and antiviral activity.

The loading of the BMSC plates and melt-blown cloth with silver and copper ions was confirmed by XRD, EDS, and XAFS experiments (Fig. 3e–h, Fig. S1) (*29–31*). Based on virus detection experiments and bacteria elimination studies (Fig. 2(c-d) and Supplementary Report 1), F-MAX shows excellent antibacterial and antiviral performance. F-MAX inactivates up to 99.99% of influenza A virus, which is mainly due to the electrostatically charged filter cartridge in the middle layer. Moreover, after 24 h, 48 h, and two weeks of operation, F-MAX shows bacterial eradication efficiencies of 99%, 98%, and 94%, respectively. These rates far exceed the sterilization efficiency of Nippon’s antiviral coating. F-MAX can be operated non-stop, providing sustained and superior bactericidal performance. This is primarily attributed to the magnesium hydroxide layer of F-MAX and the efficiently loaded silver and copper elements. Extensive bacterial studies have revealed the multiple antimicrobial mechanisms of copper, including (1) plasma membrane permeabilization, (2) membrane lipid peroxidation, (3) protein alteration, (4) inhibition of protein assembly and activity, and (5) denaturation of nucleic acids (*32*). Membrane disruption may occur due to the electrostatic force exerted by copper ions on the extracellular plasma membranes of bacteria. Protein damage occurs through the displacement of essential metals from their native binding sites on proteins or by direct interaction with proteins. Copper binding sites on nucleic acids also denature proteins. In addition, the cyclic redox reaction between Cu^+^ and Cu^2+^ is known to produce highly reactive hydroxyl radicals. Reactive oxygen species (ROS) cause or contribute to cell death through their interaction with cell membranes. Copper nanoparticles (CuNPs) have been shown to have excellent antimicrobial and antiviral surface activity due to their small size and high surface-to-volume ratio (*33–41*).

Silver is another antiviral material that inactivates viruses by interacting with viral envelopes and viral surface proteins, blocking viral penetration into cells, blocking cellular pathways, interacting with viral genomes, and interacting with viral replication factors. Silver ions can act as an early antiviral drug to disrupt viral replication. Moreover, silver also shows an inhibitory effect in the later stages of viral replication (*42–49*).

### Durability of F-MAX

The F-MAX filter cartridge exhibits remarkable filtration efficiency, which is bolstered by its robust structural integrity and layer reusability. These features provide the filter cartridge with exceptional durability and potential for recycling, setting a new standard in air purification technology.

#### Virus and Bacteria Reduction Rates

As shown in Fig. 5(a), F-MAX exhibits strong antiviral capabilities over a period of 336 h (two weeks). Initially, the virus reduction rate is very high, and this rate only slightly diminishes over time. Thus, F-MAX exhibits good long-term reliability and potential for extended use. The antibacterial performance of F-MAX is displayed in Fig. 5(b). Initially, F-MAX exhibits an impressively high bacterial reduction efficiency, and performance only slightly declines during long-term operation.

**Fig. 5.**
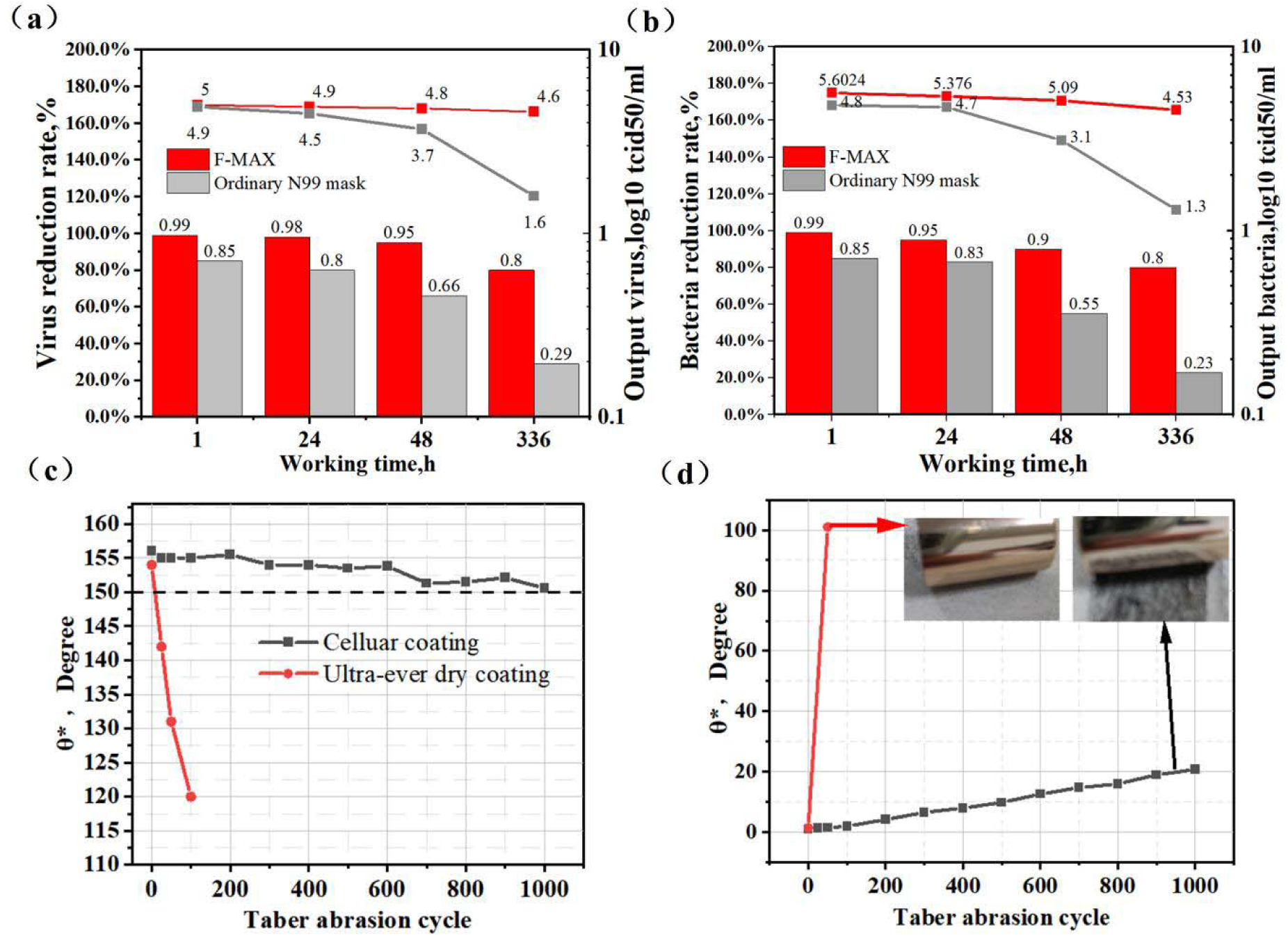
Durability of F-MAX. Overall (a) antiviral efficacy and (b) antibacterial efficacy of F-MAX and an N99 mask over time; (c)-(d) Water contact angle of cellular coating vs. Tabar abrasion cycle number.

Longevity of Self-healing Cellular Coating Under Abrasion:

Figures 5 (c) and (d) reveal the resilience of the cellular coating of F-MAX when subjected to Taber abrasion cycles. Remarkably, the water contact angle θ remains consistently high across 1000 abrasion cycles. This indicates the strongly durable hydrophobicity of the layer. Apart from this, the micro structure of the cellular The design mimics the microstructure of lotus leaves and employs composite microparticle material technology, endowing it with self-healing capabilities, thereby significantly enhancing its lifespan.demonstrating the superior construction and sustained performance of F-MAX (*50*).

##### Maintenance of Electrostatic Charge Over Time

F-MAX can retain an electrostatic charge over time, which is crucial for maintaining its ability to capture particles from the air. As shown in Fig. 4 (c) and (d), the electrostatic properties of this filter are preserved over long-term use, showcasing the enduring functionality of the cellular coating.

The DR coatings of F-MAX also display good hydrophobicity. The wear-resistant nature of the BMSC sheets means that after dust saturation, F-MAX can be readily cleansed by washing or wiping. This significantly facilitates easy reuse. The overall durability, usability, and long lifespan of F-MAX compare favorably to conventional fiberglass cloth filters. Therefore, F-MAX is an innovative and sustainable air filtration solution.

#### Carbon footprint and carbon reduction pathways of F-MAX

The composite multilayer plate filtration system reported herein represents a significant advancement in reducing the carbon emissions associated with consumable materials when compared to traditional filters. Carbon footprint (*51, 52*) calculations of F-MAX and other filtration systems are provided in Table 2 and Fig. 6. This muti-layer plate achieves a remarkable carbon emission reduction of up to 55% compared to common commercially available filters that offer the same level of air filtration efficiency. Additionally, the carbon emissions per unit of dust capacity are also reduced by up to 53%. This outstanding achievement is primarily attributed to the highly effective adsorption of larger dust particles by the two ultra-low carbon BMSC panels within this filter.

**Fig. 6.**
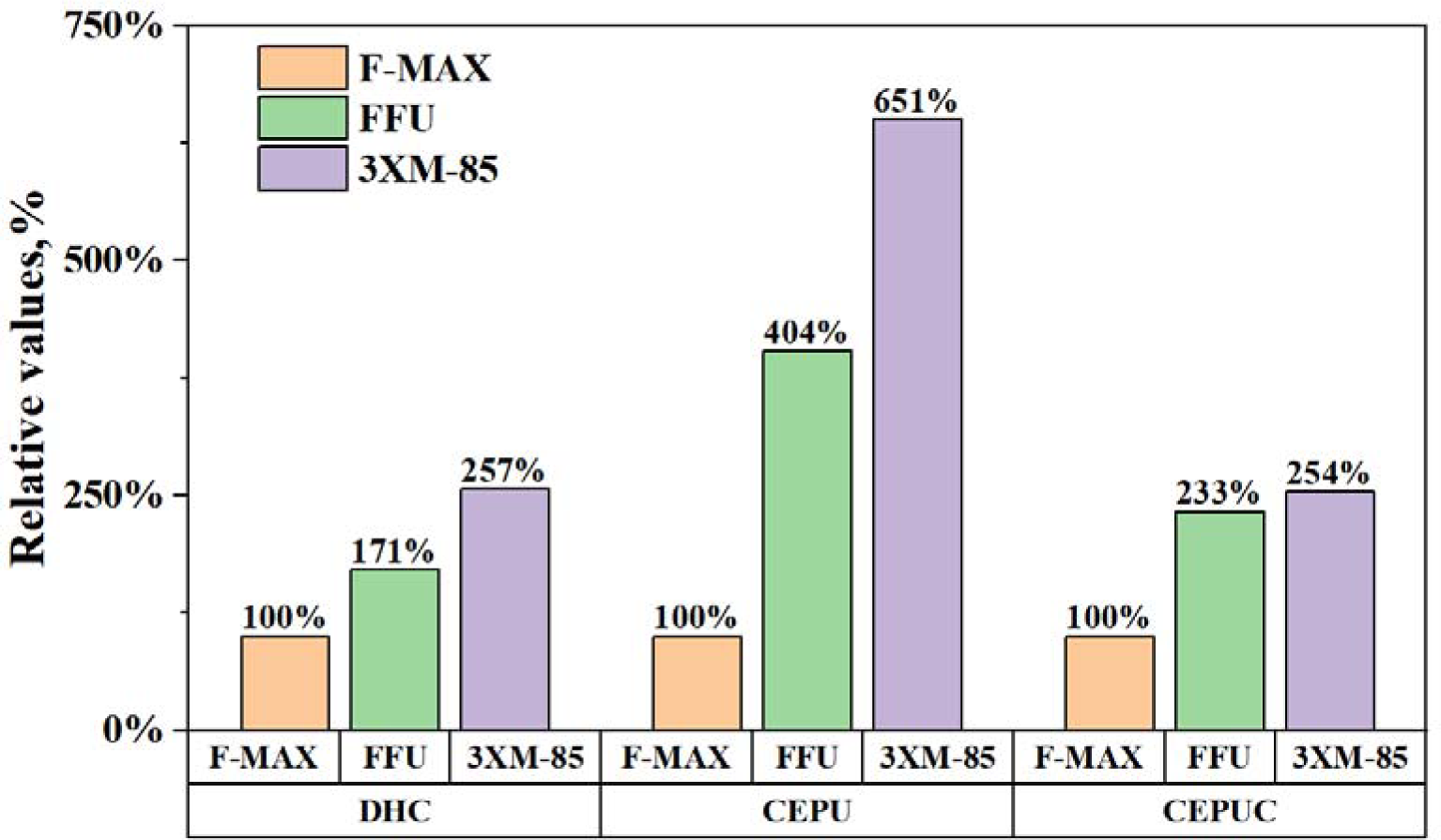
Comparison of F-MAX, FFU, and 3XM-85 carbon emissions. Besides, compared with traditional materials used to produce the filter media in other air filters, the production of BMSC results in extremely low (near-zero) carbon emissions, which is due to the following reasons:

**Table 2.** Carbon emissions of F-MAX, FFU, and 3XM-85.

| Category of filter | Filtration efficiency level | DHC | Carbon emissions Filter media | CEPU.kg/m<sup>2</sup> | Carbon emissions of filter, kg | CEPUC, g (CO<sub>2</sub>)/g (dust) |
| --- | --- | --- | --- | --- | --- | --- |
| Multilayer plate | Europe, F8,99.99% | 35 | BMSC+melt-blown+fabric + biomimetic coating | 768 | 71 | 2.02 |
| Fan filter unit (FFU) produced By Zijing Co. | Europe, F8,99.99% | 60 | Glass fiber | 9106 | 2558 | 42.6 |
| 3XM-85 Hi-Flo produced by Camfil Co. | Europe, F8,99.99% | 90 | Glass fiber | 7006 | 2552 | 28.3 |

\* DHC: Dust holding capacity per unit of filtration area.
CEPU: Carbon emissions per unit of filtration area.
CEPUC: Carbon emissions per unit of dust capacity.

1. Substituting LBM with lithium brine by-product magnesia (Li-MgO): The western lithium brine industry generates millions of tons of magnesia waste, which is mixed with impurities such as lithium and boron. This waste can be used to produce high-quality magnesia (Li-MgO) via low-temperature calcination (700–750 °C). Recent research by our team has demonstrated that Li-MgO exhibits superior hydration activity compared to LBM. BMSC materials produced using Li-MgO perform nearly as well as those made with LBM, so the performance of filter media is not compromised. Therefore, the substitution of LBM with Li-MgO is both feasible and offers promising environmental benefits.The carbon emissions per ton associated with the production of Li-MgO are only 9% of those generated by LBM, and there are two main reasons for this. One, calcination temperatures in the range of 700–750 °C are required for Li-MgO. This is significantly lower than the 850–1000 °C required for LBM production, resulting in substantial energy savings. Two, the primary component of raw LBM is magnesium carbonate, whose calcination releases a substantial amount of carbon dioxide (1.4 tons of CO_2_ per ton of magnesia). In contrast, the raw material used for the production of Li-MgO production is a mixture of magnesium oxide and magnesium hydroxide. Calcining this mixture does not produce (0 tons of CO_2_ per ton of magnesia).
2. Substituting sulfuric acid-derived magnesium sulfate with flue gas desulfurization by-product sulfuric acid: Traditionally, magnesium sulfate is primarily produced by reacting magnesium oxide with sulfuric acid, resulting in significant carbon emissions (indirect production of 0.4 tons of carbon dioxide per ton of magnesium sulfate). In contrast, the utilization of magnesium in flue gas desulfurization produces magnesium sulfate as a by-product in a zero-carbon-emission process. In this work, the use of this by-product magnesium sulfate, which is required for BMSC production, was investigated. According to the results, BMSC prepared using industrial flue gas desulfurization by-product magnesium sulfate exhibits the same permeability and basic physical properties as traditional BMSC. Therefore, the use of magnesium sulfate produced as a by-product during flue gas desulfurization instead of traditionally prepared magnesium sulfate is entirely feasible for the production of BMSC.

Therefore, the simultaneous use of Li-MgO and industrial flue gas desulfurization by-product magnesium sulfate in the production of ultra-low-carbon BMSC (ULC-BMSC) can significantly reduce overall carbon emissions.

The low-carbon nature of the prepared multilayer plate can mainly be attributed to the use of BMSC.

#### Design of F-robot and its Application

The newly developed F-robot, a state-of-the-art multi-layer composite air purifier showcased in the attached images, represents a leap forward in air purification technology, specifically engineered for the demanding environment of laboratory animal rooms. It is tasked with the critical role of eliminating viruses, bacteria, odors, and fine particulates, thus ensuring a pristine air quality that meets rigorous laboratory standards. The purifier’s sophisticated filtration system, as illustrated in Fig. 7(b), comprises a series of expertly crafted layers: a uniformly fine foamed basic magnesium sulfate cement paste (FBMSCP), a robust ultra-durable coating, a long-lasting electrostatically charged melt-blown filter material, and a hydrophobic Desert Rose coating. Together, these layers form a formidable barrier, capturing particles down to 0.003 microns and eliminating 99.9% of harmful microorganisms.

**Fig. 7.**
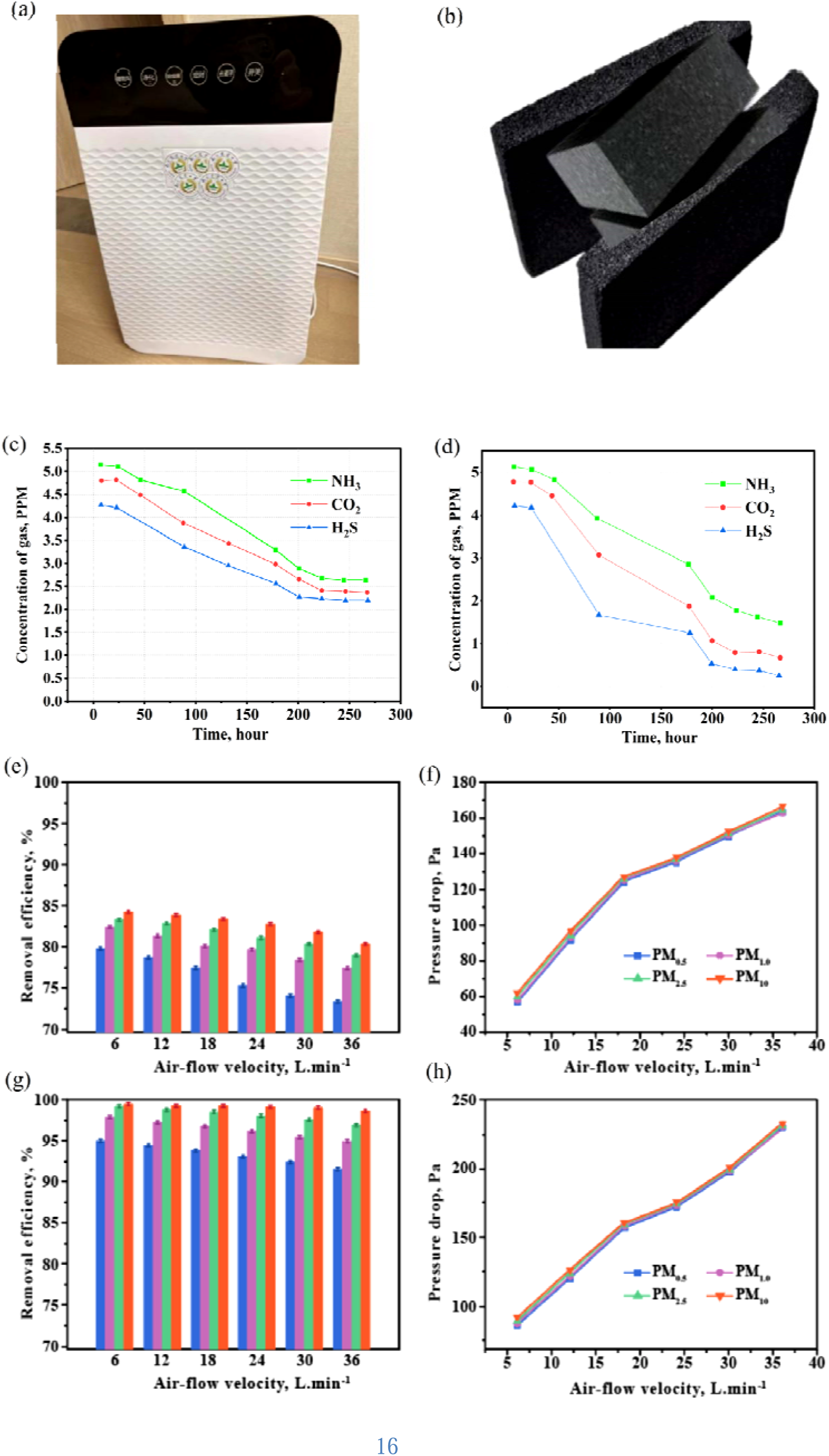
Overall structure and function of F-robot: (a) The finally product of F-robot, (b) F-MAX after function, (c) The ability of reduce pollutants based on commercial air purifier machine in mouse room. (d) The ability of reduce pollutants based on F-robot in mouse room. The pollutants here mean ammonia (NH_3_), carbon dioxide (CO_2_), and hydrogen sulfide (H_2_S), (e–f) Removal efficiency and pressure drop of commercial air purifier machine in mouse room. (g–h) Removal efficiency and pressure drop of F-robot in mouse room. (blue, purple, green and orange correspond to PM0.18, PM0.3, PM2.5, and PM10), respectively.

The effectiveness and long residual action of F-robot is further underscored by the graphical data provided in images Fig. 7(d). Compare the effectiveness of commercial air purifier machine Fig. 7(c), the F-robot shows evident advantage. It illustrates the superior performance of this cartridge. A significant reduction in the concentrations of ammonia (NH_3_), carbon dioxide (CO_2_), and hydrogen sulfide (H_2_S) can be observed within 300 h. The initial phase of the F-robot is characterized by a notable decrease in the levels of CO_2_ and H_2_S, indicating its robust and rapid absorption properties. While the concentration of NH_3_ declines more gradually, this trend demonstrates the long-term efficacy of the filter, which is an essential feature for scenarios requiring durable filtration solutions. The error bars demonstrate the reliability of this filtration process, with minimal fluctuations observed. Therefore, the cartridge has good long-term consistency. Moreover, no rapid saturation phenomenon can be observed. These results demonstrate that F-robot is an ideal candidate for environments consistently exposed to noxious gases. By the way, the actual use of which was also shown in Fig. S3, which depict the concentration of various gases over time.

Besides, as illustrated in Fig. 7(e), the removal efficiencies of PM based on commercial air purifier machine are less than 83%. And they endow it characteristics of high pressure drop (Fig. 7(f)). While the removal efficiencies of PM based on F-robot shows evident higher removal efficiency (Fig. 7(g)), but also a higher pressure drop (Fig. 7(h)). Besides, the the removal efficiencies of PM based on F-robot illustrates the superior performance. A significant reduction in the concentrations of the removal efficiencies of PM based on ordinary commercial filter cartridge can be observed within 36 min.

A prominent feature of the F-robot is the chromatic sensor embedded within its design, as seen in Fig. 7(a). This sensor offers an immediate readout of the air quality index, allowing for a visual comprehension of the ambient air conditions. The purifier’s advanced sensing technology delivers real-time air quality monitoring, as indicated by the clear display panel, enabling consistent awareness and peace of mind regarding the air status.

The F-robot employs a multifaceted approach to pathogen removal, utilizing silver ion and copper element-based sterilization along with electrostatic adsorption—a testament to the purifier’s rigorous design philosophy aimed at creating a sterile environment. Its 460° circular airflow design, distributes purified air evenly, enhancing the overall efficacy of the system.

With its sleek appearance and discreet operational profile, the F-robot is not only a powerhouse of purification but also an aesthetically pleasing addition to any laboratory setting. The purifier’s environmental credentials are equally impressive, with sustainable material construction and energy-efficient performance aligning with eco-friendly practices.

The intelligent controls of the F-robot, intuitive to use and featured on the purifier’s facade, make it user-friendly and accessible. Users can track air quality trends, manage settings remotely, and stay updated with maintenance needs, ensuring the purifier operates at optimal levels at all times.

In sum, the F-robot, through its innovative design, documented performance, and user-centric features, encapsulated in the provided images, is a paradigm of modern air purification solutions.

It not only meets the immediate demands for clean air in sensitive environments but also sets a precedent for future developments in the field, aiming for a healthier, cleaner tomorrow.

## Discussion

The F-MAX multilayer composite plate is at the forefront of air filtration technology, combining high efficiency with an environmentally friendly design, marking a significant step forward in aligning air purification with environmental conservation. Its advanced multilayer structure captures a broad range of pollutants, with each layer tailored to target specific particles, ensuring thorough filtration. The F-MAX’s high filtration efficiency is unmatched, capable of filtering air contaminants with exceptional precision and significantly enhancing air quality. This system also boasts antiviral and antibacterial properties, proactively preventing the spread of pathogens, crucial in today’s world where airborne disease transmission is a persistent public health challenge.

The durability of F-MAX is paramount, designed to withstand long-term use without compromising performance, making it a sustainable choice for air purification needs. In terms of environmental impact, F-MAX contributes to carbon emission reduction and features eco-friendly recyclability, lessening the need for frequent filter replacements and supporting global efforts to reduce environmental footprints. The versatility of F-MAX makes it suitable for both commercial and residential applications, capable of adapting to various environments to ensure healthier living and working spaces.

In essence, the F-MAX multilayer composite plate transcends being just a product; it represents a transformative move towards healthier living environments, adeptly removing harmful particles and pathogens from the air, boasting a long lifespan, and embodying environmental responsibility. With its introduction to the market, F-MAX signifies not just a technological advancement but also a dedicated commitment to a sustainable future. As it gains recognition across sectors, its anticipated influence on indoor air quality and ecological practices is poised to be significant, establishing F-MAX as a crucial element in the pursuit of a cleaner, safer, and greener future.

### Newly developed and used technology of F-robot

#### Novel fine foaming technology tailored for magnesium cement

This study introduces a novel fine foaming technology tailored for magnesium cement, traditionally used as a construction material. We have developed a new foaming agent and a method that optimizes the basic components, foaming, stabilizing, and regulating elements. These improvements significantly enhance the foaming capability of the agent and the stability and fineness of the foam. Additionally, we have innovatively optimized the unevenness of the foam pores in traditional foamed magnesia panels. The conventional magnesium foam mortar often has inconsistent pore sizes, with large voids unable to effectively filter air, and the mortar interfaces around these voids being too dense for air passage. However, our newly developed foamed magnesia panels feature uniformly fine pores. (Comparison between traditional and newly developed foaming magnesium cement are shown in following Fig. S4) The pore and airway diameters are adapted to the crystalline whiskers of the 5-1-7 basic magnesium sulfate, achieving optimal filtration efficiency. The development of this technology not only expands the application range of magnesium cement but also opens new possibilities for manufacturing more efficient air filtration materials.

#### Novel electrostatic electret technology

Drawing from the core technology of gel polymer hydrogel electrolytes (GPHEs) featuring graphene oxide (GO) and polyvinyl alcohol (PVA) (*53, 54*), we developed a novel, washable melt-blown fabric filter core (Fig. 3(d)). This filter core could integrate the exceptional ion conductivity and self-healing properties of GPHEs to enhance filtration without sacrificing breathability.

The newly developed melt-blown fabric filter core would leverage the high ion conductivity of GPHEs, allowing it to attract and capture airborne particles effectively. The soaking process in a ZnSO_4_/MnSO_4_ solution, as used in GPHEs, could be adapted to imbue the fabric with electrostatic properties, enhancing its particle capture efficiency without a corresponding increase in air resistance.

Moreover, the filter core would inherit the GPHEs’ remarkable stretchability and durability, making it suitable for reusable and long-lasting air filtration applications. The low Young’s modulus of the GPHEs suggests that the filter core would be flexible, which is an advantageous property for installation and maintenance in various air purifying devices.

To address the challenge of maintaining filtration efficiency over time, the filter core could incorporate the self-healing characteristics of GPHEs. This would allow it to retain its filtration capabilities even after multiple washing cycles, reducing the need for frequent replacements and thus lessening the environmental impact.

By incorporating graphene oxide and PVA hydrogel matrix technology into the melt-blown fabric, the filter core would not only exhibit high filtration efficiency and stability but also demonstrate low production costs and a high energy density, translating into an economical and energy-efficient solution for air purification.

Such a filter core, with its high filtration efficiency, stretchability, self-healing properties, and low cost, offers an innovative solution that aligns with the sustainability goals of reducing waste and promoting recyclable materials in air purification technology.

#### Combined used structure of F-MAX

Inspired by biomimetic multilayer structures, a multilayered composite filter was designed to reduce the presence of harmful viruses, bacteria, and dust in air. This plate consists of 5 layers, as shown in Fig. 1. Two layers are BMSC plates reinforced by a glass fiber net. These panels have a thickness of 2.5 cm and are each covered with a bionic antiviral coating. These coatings are used to cover both sides of the filter plate. The entire system is glued together using MPC. The center of the structure contains an electrostatically charged melt-blown cloth capable of maintaining a suitable electrostatic load. The potential for using this multilayer plate in building structures was evaluated by measuring its mechanical properties. The experimental results demonstrate that this plate can fulfill the strength requirements of both national and international standards. FBMSCP exhibits compressive and splitting tensile strengths of more than 8.5 MPa and 1.2 MPa, respectively. Moreover, the overall average splitting tensile strength of the plate is

1.8 MPa, which is significantly higher than that required by the relevant standards. Therefore, the structure of this plate exhibits superb stability and durability. These results indicate that this plate will potentially have a longer working lifetime compared to ordinary filters, and it can independently function without the physical support of an additional framework structure. The specific materials of each layer in the filter plate are shown in Supplementary Materials.

## Acknowledgments

Many thanks to Professor Pengfeng Xiao’s research group for providing support on the knowledge and preparation methods related to low-carbon alkaline magnesium cement mortar.

## Author contributions

Conceptualization: Zhongjie Fei, Ping Jiang, Haotian Fan, Xing Liu

Methodology: Zhongjie Fei, Haotian Fan, Pingjiang

Investigation: Zhongjie Fei, Haotian Fan,Xingliu, Pengfeng Xiao

Visualization: Haotian Fan, Zhongjie Fei, Ping Jiang,Xing Liu

Funding acquisition: Zhongjie Fei, Ping Jiang, Xing Liu, Pengfeng Xiao

Project administration: Zhongjie Fei, Ping Jiang, Xing Liu, Pengfeng Xiao

Supervision: Liujun Xiao, Mengjie Li, Song Ge

Writing – original draft: Zhongjie Fei, Haotian Fan

Writing – review & editing: Zhongjie Fei,

## Data and materials availability

All data are available in the main text or the supplementary materials.

## Supplementary Materials

### Materials

#### Desert Rose coatings

The BMSC plate’s second layer is coated with a “Desert Rose” biomimetic material, mimicking the hydrophobic traits of desert plants. Its rose-like microscopic structure and organic substrate grant it superior hydrophobicity, effectively repelling water and attracting nanoscale impurities. The coating’s layered, rose petal-like morphology provides a robust adsorption force, ensuring high filtration efficiency.

Experimental data using contact angle measurements before and after coating. Foam without coatings show hydrophilic while it coated by cellular coatings and reveal significant hydrophobic enhancement. The contact angle of foam without coatings is 19° while that of coated by cellular coatings is 125°.

reveal significant hydrophobic enhancement. Initially hydrophilic (shown as Figure S5), the surface becomes water-repellent post-coating, evident from the increased contact angle. This change highlights the coating’s ability to modify surface properties, enhancing water resistance.

The Desert Rose’s porous structure ensures near-complete particle capture during filtration. Its unique needle-like form maintains over 95% filtration efficiency for sub-micron particles. Remarkably, the bionic filter sheet, washable and reusable, maintains its efficiency beyond five washes, making it an eco-friendly and cost-effective solution for the final layer in the F-MAX system.

#### Foaming basic magnesium sulfate cement paste

Building upon Professor Yu Hongfa’s(*55*) alkaline magnesium sulfate cement technology, a mixture of magnesium sulfate heptahydrate, lightly calcined reactive magnesium oxide, fine aggregates, a carboxylate admixture, a foaming agent, and water were used in precise proportions to modify basic magnesium sulfate cement (BMSC) with 5Mg(OH)_2_ · MgSO_4_ · 7H_2_O (5 · 1 · 7 phase) crystalline whiskers(*56*). Compared to traditional 3·1·8 phase magnesium sulfate cement, the modified 5·1·7 phase BMSC offers several advantages, including rapid setting, early strength development, high strength, good water resistance, corrosion resistance, and carbonation resistance (the physical characteristics of this modified BMSC are reported in the Supplementary Text). The mechanical properties, reinforcement performance, and durability of 5·1·7 phase BMSC are comparable to those of Portland cement, and this material even surpasses ordinary Portland cement in some aspects(*57–58*). Thus, this BMSC material is a highly reliable new type of cementitious material. Due to the high strength of the 5·1·7 phase crystalline whiskers (Figure 3(d)) as well as the early strength (Table S1) and high durability of BMSC (which grows in a needle-like manner along pore structures), fine pores with excellent structural stability can be formed under the action of foaming agents. When used as a multi-pore filter core, this material can efficiently filter fine particulate matter from air. Therefore, 5·1·7 phase BMSC with crystalline whiskers is a suitable material for the outer layer of the filter plate. The specific mortar formula and corresponding physical performance parameters are reported in the table below. It should be noted that except for the admixture, all the raw materials used in this board are industrial by-products and waste materials.

#### F-MAX electrostatically charged filter cartridge in the middle layer

The primary constituents of the filter cartridge are polypropylene fibers with diameters ranging from 0.6 to 1.5 µm, as depicted in Figure 3(c). Employing a distinctive approach, silver ions are integrated into the melt-blown fabric to provide antibacterial and antiviral properties. The melt-blown fabric is prepared using a blend of polypropylene, electrolytic stone, silica, silver ions, and copper ions. This fabric undergoes electrostatic polarization, which significantly enhances its charge and provides antiviral capabilities. Due to the combination of inorganic metals and non-metallic elements, the melt-blown fabric provides highly efficient and long-term filtration and antiviral functionalities.

A symmetric layer structure is adopted for the multilayer composite plate designed in this study, enabling its utility as a construction material. The design of which was conducted on the base of systematical analysis and ML developed through the Computer simulation data (Figure S6). In addition to its impressive filtration and antiviral properties, this plate exhibits superb structural stability, surpassing the standards of conventional prefabricated foam concrete boards.

Crucially, the optimal blend of electrolytic stone and polypropylene fibers coupled with the inclusion of high-capacitance silver and copper ions provides the cartridge with exceptional electrostatic storage capacity and durability. Furthermore, the detachable design of the F-robot facilitates the repetitive and efficient use of the F-MAX electrostatically charged filter cartridge through advanced polarization. This prolongs the lifespan of F-MAX and enhances its sustainability. Thus, an overall reduction in carbon emissions is achieved compared with traditional filtration systems.

#### Ultra-durable superhydrophobic cellular coatings

To filter the passing air in a hierarchical manner, ultra-durable superhydrophobic cellular coatings were selected for the final layer of the F-MAX filter cartridge. Due to their special structure, these monocellular coatings(*59*) provide superhydrophobicity, multiphase repulsion, and ultra-long-term performance. Each cell is composed of a hard porous diatomaceous earth microshell and a releasable nanoseed, which has unique physicochemical properties(*60–63*). In the coating, cake-shaped cells and partially fragmented nanoseeds can be observed. A “self-healing” function is achieved under large loads when the nanoseeds released by the unit cells are broken, which maintains the ultra-long-term corrosion resistance and hydrophobicity of the coating. There is a microporous gap of less than 10 µm between the cells of the coating, which enables the initial filtration of coarse particles without absorbing the water from the filtered air.

## Supplementary Materials

### Materials

#### Desert Rose coatings

Experimental data using contact angle measurements before and after coating reveal significant hydrophobic enhancement. Initially hydrophilic (shown as Figure S5), the surface becomes water-repellent post-coating, evident from the increased contact angle. This change highlights the coating’s ability to modify surface properties, enhancing water resistance.

#### F-MAX electrostatically charged filter cartridge in the middle layer

A symmetric layer structure is adopted for the multilayer composite plate designed in this study, enabling its utility as a construction material. The design of which was conducted on the base of systematical analysis and ML developed through the withdrawn experimental data,(Figure S6) In addition to its impressive filtration and antiviral properties, this plate exhibits superb structural stability, surpassing the standards of conventional prefabricated foam concrete boards.

Crucially, the optimal blend of electrolytic stone and polypropylene fibers coupled with the inclusion of high-capacitance silver and copper ions provides the cartridge with exceptional electrostatic storage capacity and durability. Furthermore, the detachable design of the T-robot facilitates the repetitive and efficient use of the F-MAX electrostatically charged filter cartridge through advanced polarization. This prolongs the lifespan of F-MAX and enhances its sustainability. Thus, an overall reduction in carbon emissions is achieved compared with traditional filtration systems.

#### Ultra-durable superhydrophobic cellular coatings

To filter the passing air in a hierarchical manner, ultra-durable superhydrophobic cellular coatings were selected for the final layer of the F-MAX filter cartridge. Due to their special structure, these monocellular coatings (*59*) provide superhydrophobicity, multiphase repulsion, and ultra-long-term performance. Each cell is composed of a hard porous diatomaceous earth microshell and a releasable nanoseed, which has unique physicochemical properties(*60–63*). In the coating, cake-shaped cells and partially fragmented nanoseeds can be observed. A “self-healing” function is achieved under large loads when the nanoseeds released by the unit cells are broken, which maintains the ultra-long-term corrosion resistance and hydrophobicity of the coating. There is a microporous gap of less than 10 µm between the cells of the coating, which enables the initial filtration of coarse particles without absorbing the water from the filtered air.

Supplementary Text and report

## Supplymentary report 1: XAFS report for Cu and Ag loaded on the middle layer of F-MAX electrostatically charged filter cartridge

### XAFS data analysis report

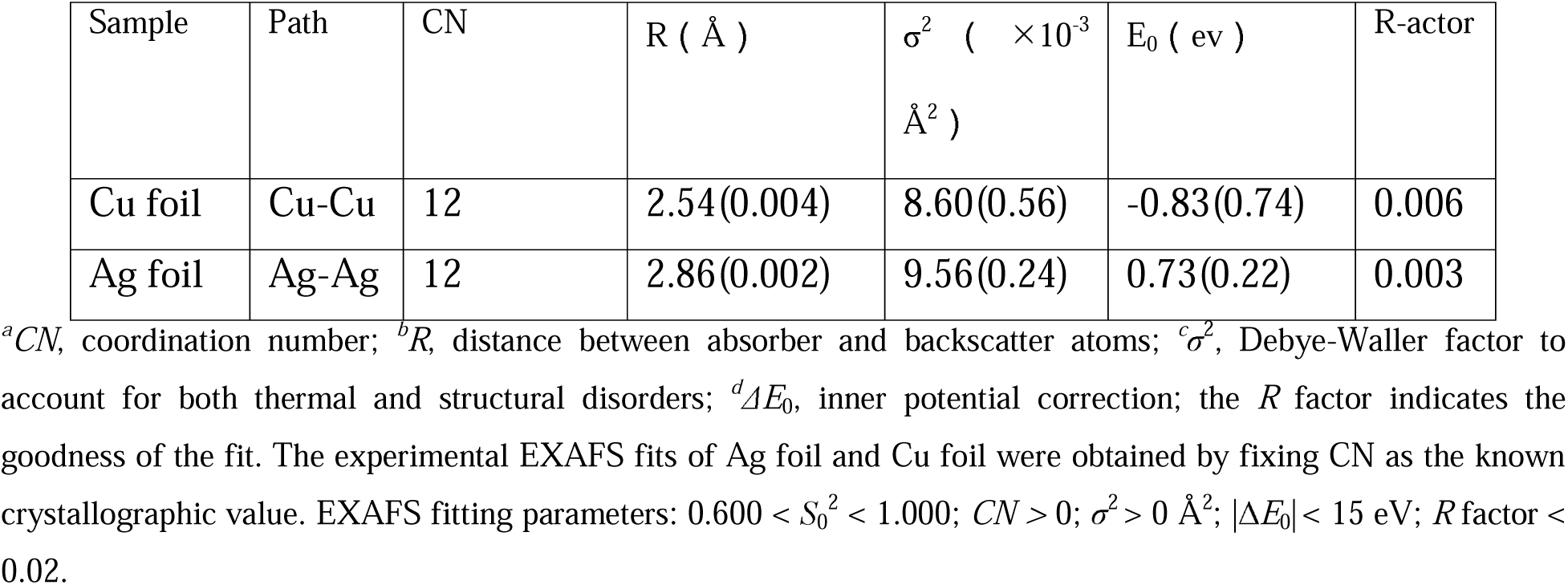
**Table** EXAFS fitting parameters at the K-edge for various samples.

1. Simple Analysis of the E&R&k Space: Analysis from the E-edge data of the elements (Figure 1) indicates that the standard Ag foil sample is in the zero-valent state, with a weaker white line peak intensity that corresponds to the actual situation.

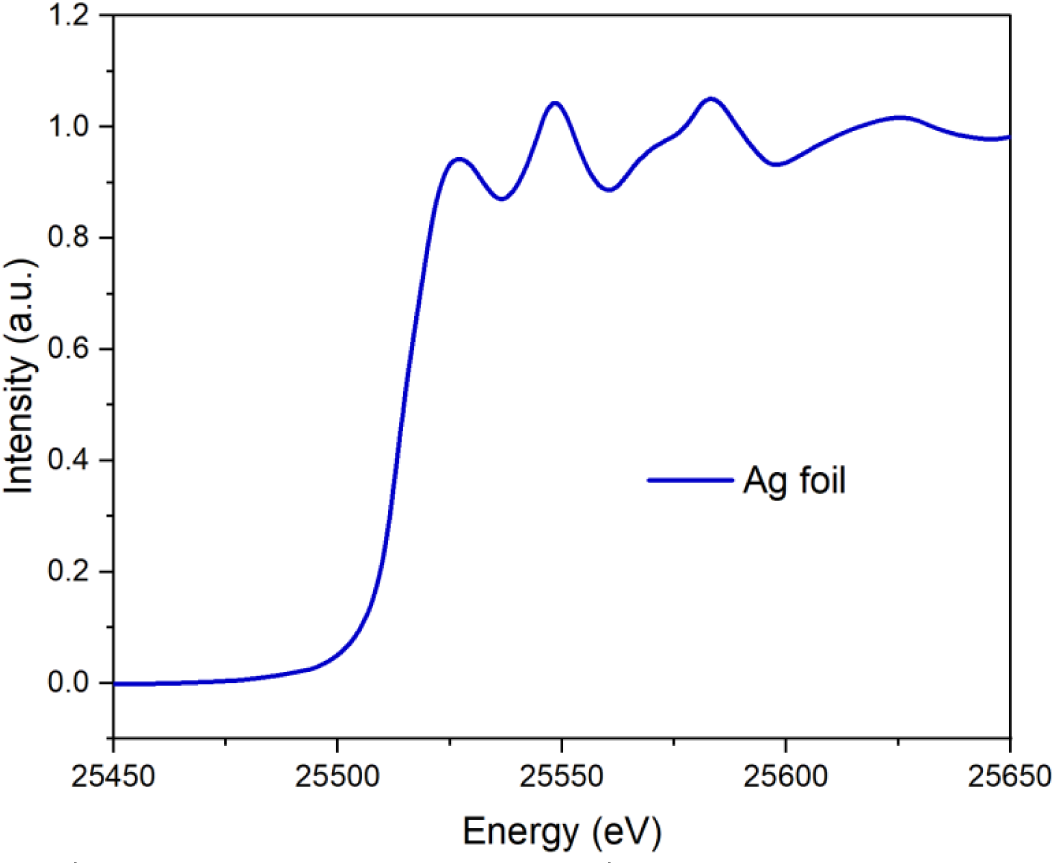 Analyzing the R-space of Ag shows that the main sample peak is in the range of 2–3 Å, which is generally ascribed to metallic Ag. Because the tested sample is elemental silver (Ag), this main peak can be attributed to Ag-Ag bonding.

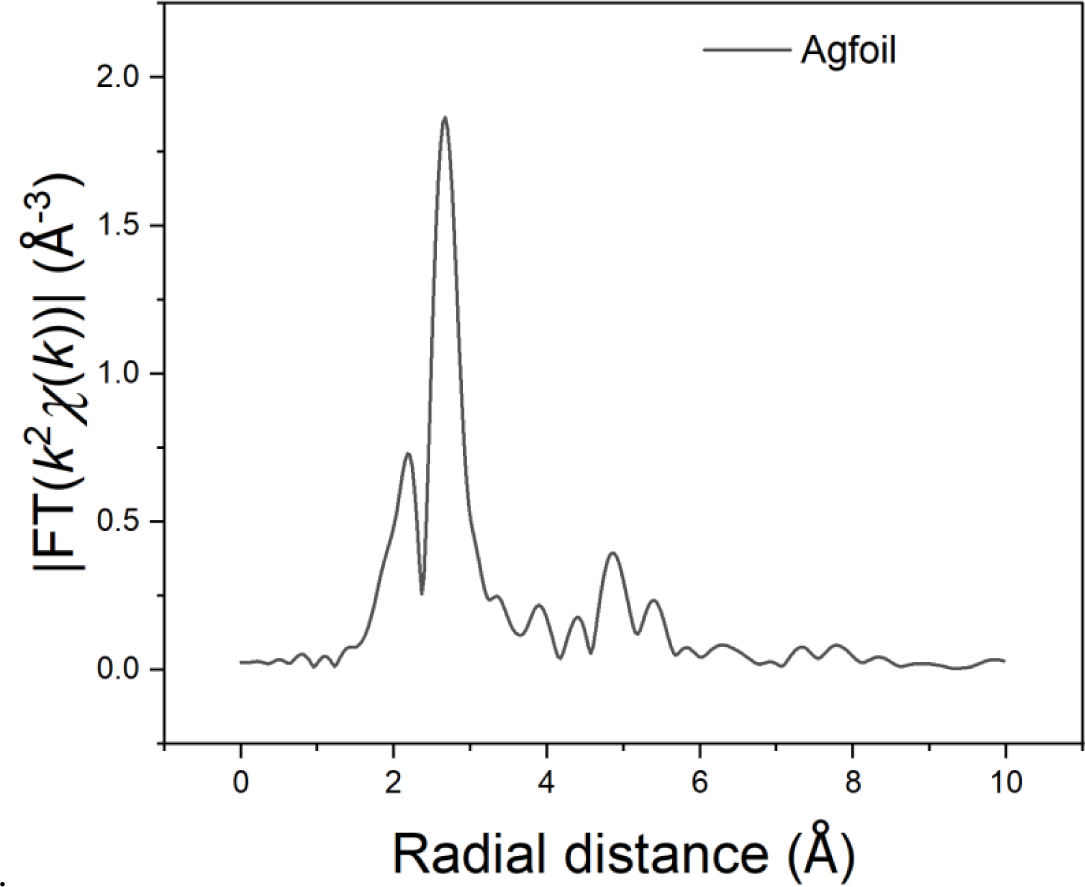
2. Fitting the sample curve to the standard structural model of elemental silver (Ag) shows a very high degree of overlap. All fitting parameters are within a reasonable range. Thus, the coordination structure present in the sample is Ag-Ag. Detailed fitting information can be found in the table above. Data reduction, data analysis, and EXAFS fitting were performed and analyzed with the Athena and Artemis programs of the Demeter data analysis packages (reference 1), which utilized the FEFF6 program (reference 3) to fit the EXAFS data. The energy calibration of the sample was conducted using a standard Ag foil, which was simultaneously measured as a reference. A linear function was subtracted from the pre-edge region. Then, the edge jump was normalized using Athena software. The χ(k) data were isolated by subtracting a smooth, third-order polynomial approximating the absorption background of an isolated atom. The k3-weighted χ(k) data were Fourier transformed after applying a Hanning window function (Δk = 1.0). For EXAFS modeling, the global amplitude EXAFS (CN, R, σ2, and ΔE0) were obtained by the nonlinear fitting (with least-squares refinement) of the EXAFS equation to the Fourier-transformed data in R-space using Artemis software.

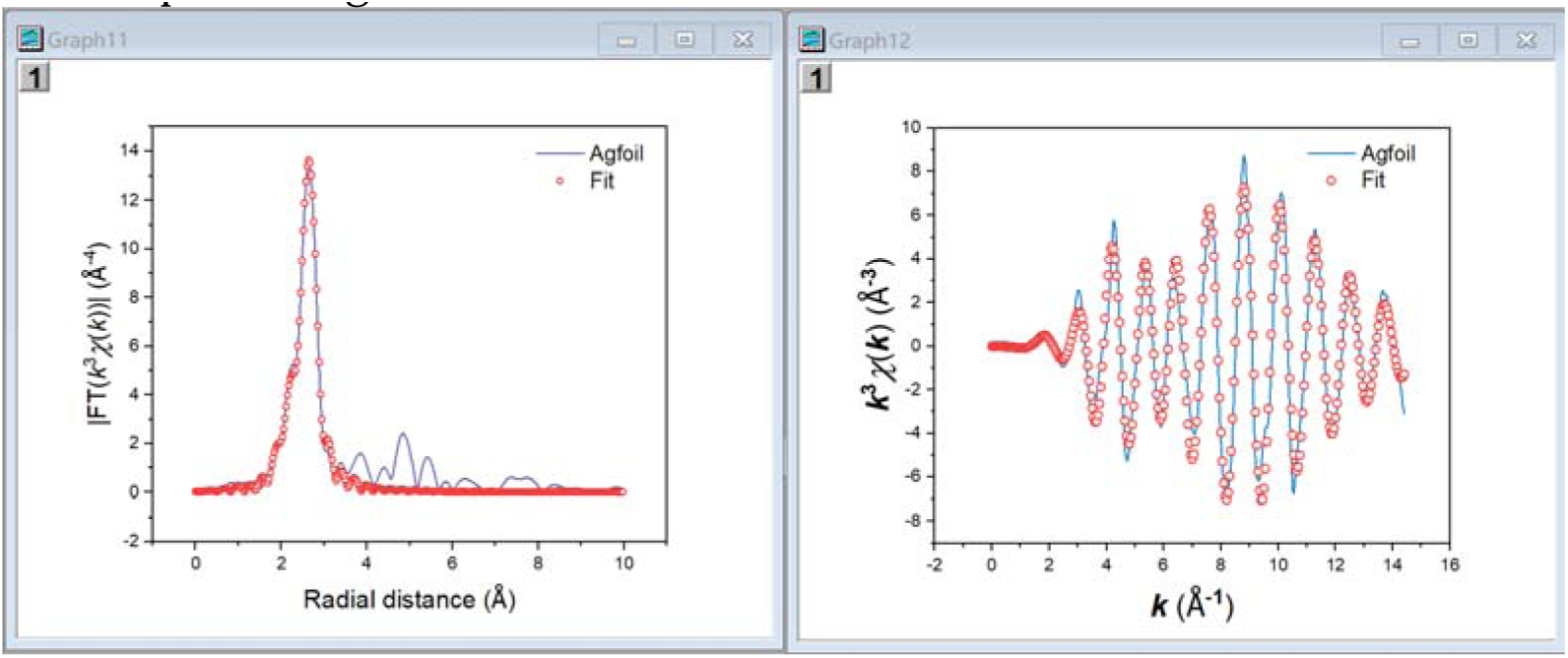
3. Wavelet Data: Different colors represent the intensity of peaks. These data indicate the distance of the coordinating atoms (i.e., bond length) and differentiate the types of coordinating atoms. A horizontal coordinate further to the right indicates a higher atomic number. In conjunction with the R-space fitting information, the wavelet data can effectively discern the coordination of the sample. The wavelet analysis shows that the main coordination in the sample is predominantly Ag-Ag.

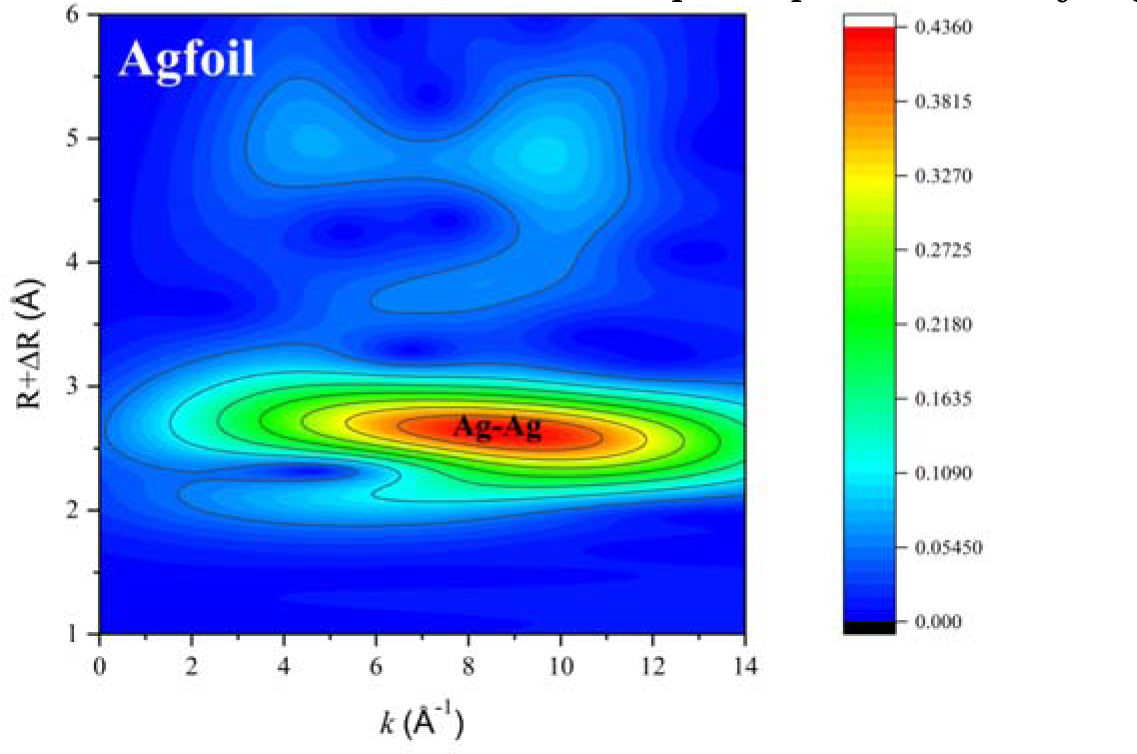

1. Simple Analysis of the E&R&k Space: Analysis from the E-edge data of the elements (Figure 1) indicates that the standard Ag foil sample is in the zero-valent state, with a weaker white line peak intensity that corresponds to the actual situation.

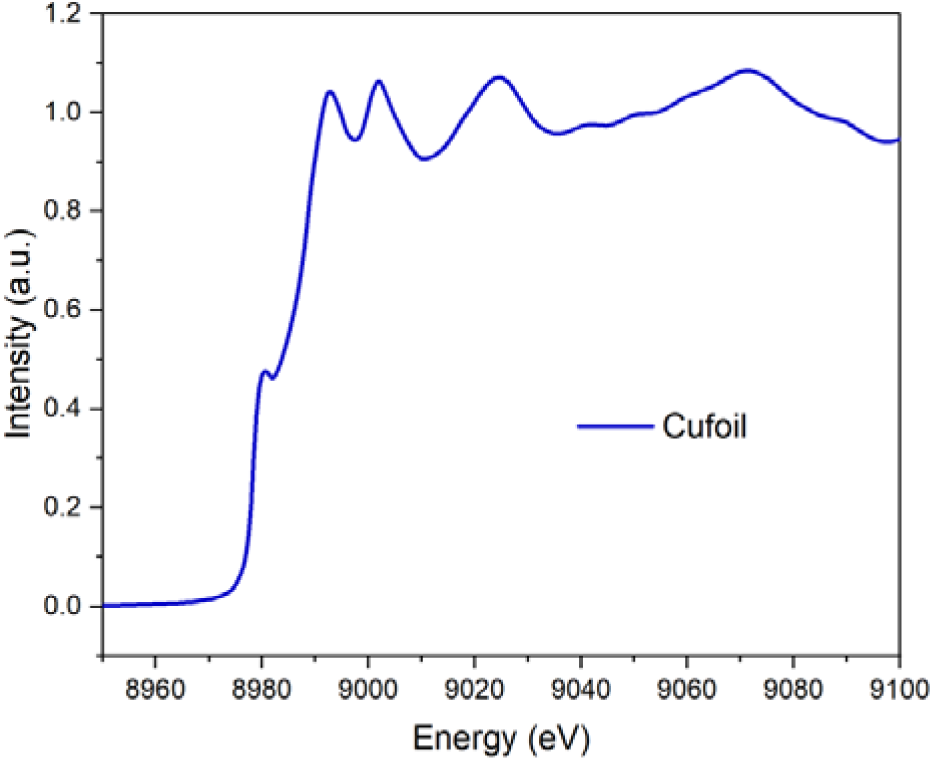 Analyzing the R-space of Cu shows that the main sample peak is in the range of 2–3 Å, which is generally ascribed to metallic Cu. Because the tested sample is elemental copper (Cu), this main peak can be attributed to Cu-Cu bonding.

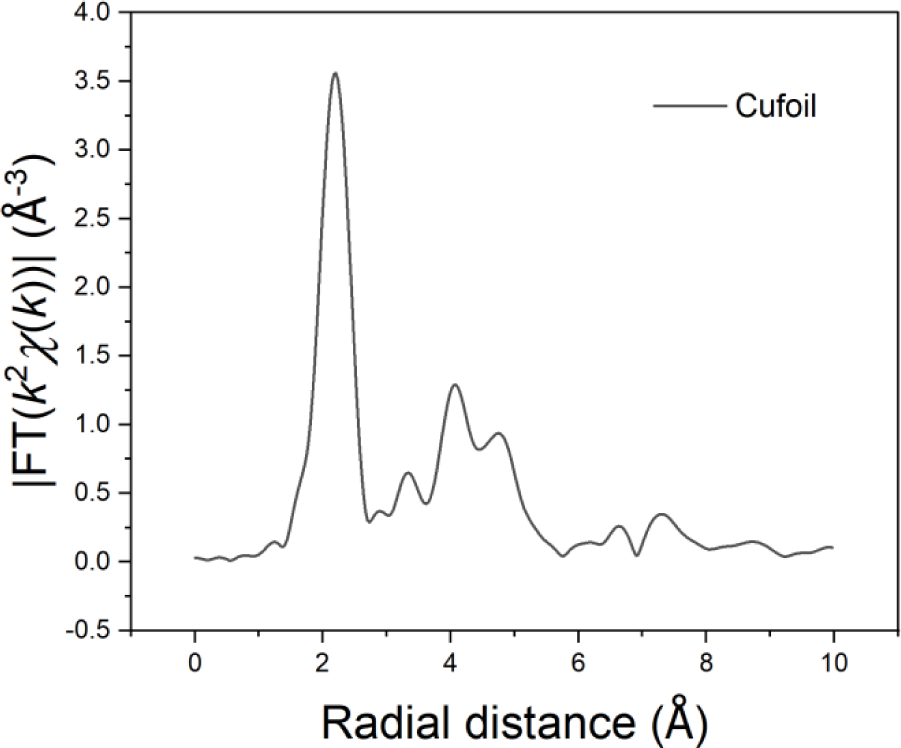
2. Fitting the sample curve to the standard structural model of elemental silver (Cu) shows a very high degree of overlap. All fitting parameters are within a reasonable range. Thus, the coordination structure present in the sample is Cu-Cu. Detailed fitting information can be found in the table above. Data reduction, data analysis, and EXAFS fitting were performed and analyzed with the Athena and Artemis programs of the Demeter data analysis packages (reference 1), which utilized the FEFF6 program (reference 3) to fit the EXAFS data. The energy calibration of the sample was conducted using a standard Cu foil, which was simultaneously measured as a reference. A linear function was subtracted from the pre-edge region. Then, the edge jump was normalized using Athena software. The χ(k) data were isolated by subtracting a smooth, third-order polynomial approximating the absorption background of an isolated atom. The k3-weighted χ(k) data were Fourier transformed after applying a Hanning window function (Δk = 1.0). For EXAFS modeling, the global amplitude EXAFS (CN, R, σ2, and ΔE0) were obtained by the nonlinear fitting (with least-squares refinement) of the EXAFS equation to the Fourier-transformed data in R-space using Artemis software.

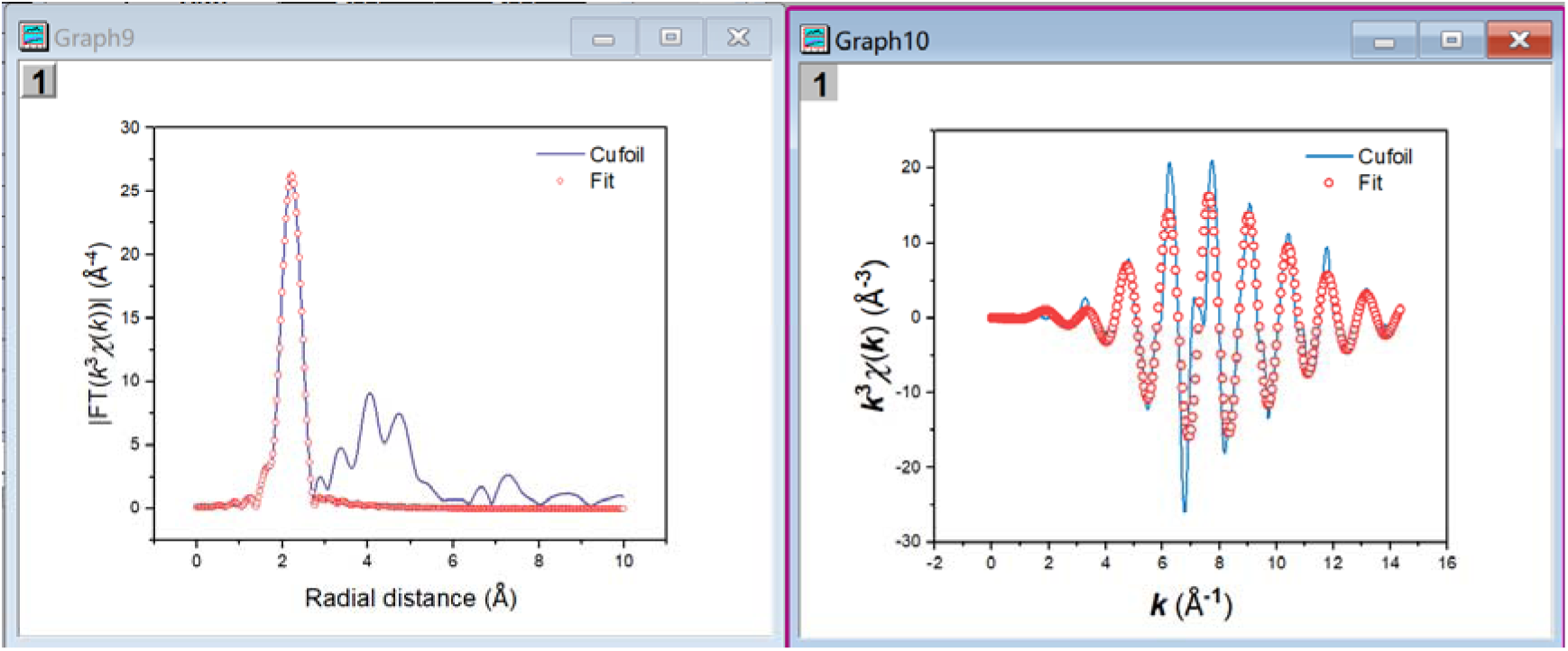
3. Wavelet Data: Different colors represent the intensity of the peaks. These data indicate the distance of the coordinating atoms (i.e., bond length) and identify the types of coordinating atoms. A horizontal coordinate further to the right indicates a higher atomic number. Combining the wavelet data with the R-space fitting information enables the coordination situation of the sample to be identified. The wavelet analysis shows that the predominant coordination in the sample is Cu-Cu.

## Supplementary report 2: Test experiment and corresponding report of antiviral efficiency of F-MAX electrostatically charged filter cartridge

### Experiment Objective

The aim of this experiment is to test the antiviral activity of a new type of virus-killing composite air filter material and verify its effectiveness in practical applications.

### Experimental Materials

Cells and Reagents: The new virus-killing composite air filter material is provided by the customer, along with Influenza Virus H1N1 (A/Puerto Rico/8/34), MDCK cells, DMEM medium, MEM medium, FBS.

#### Experimental Equipment

Includes a vertical refrigerated display cabinet, horizontal freezer, electronic balance, tabletop high-speed refrigerated centrifuge, electric constant temperature water bath, laboratory ultrapure water machine, vortex mixer, palm centrifuge, enzyme label detector, CO_2_ incubator, autoclave, clean workbench, biosafety cabinet, electronic balance, inverted microscope, etc.

#### Main Experimental Consumables

Include 1.5ml centrifuge tubes, pipettes of various sizes, etc.

### Experimental Procedures

#### Sample Preparation

Prepare the sample in aseptic culture broth and transfer the specimen aseptically into a sterile vial.

#### Sample Inoculation

Use a micropipette to accurately drop the prepared 0.2 ml virus suspension onto several points of the sample in the vial, then cover with the lid.

#### Contact Time

Incubate for 2 hours at 25°C. The contact time can be adjusted as needed but should not exceed 24 hours.

#### Virus Elution

Immediately add elution fluid to the control group test vials, vortex mix for 5 seconds, repeat 5 times to elute the virus from the sample.

#### Post-Elution Processing

After the prescribed time (2 hours), repeat the elution process as in step 4.

#### Virus Dilution

Dilute the eluted virus fluid to the eighth dilution level.

#### Cell Infection

Add the virus fluid of each dilution level to the cell plates with MDCK cells, 5 replicate wells for each dilution, and mark accordingly. Incubate the cell plates in a 37°C, 5% CO2 incubator for 3 days and observe the cytopathic effect in the wells.

### Experimental Result Analysis

#### Observation and Recording

After the experiment, observe the cytopathic effect in the wells of the cell plate and record the cytopathic wells for each dilution level.

#### Data Analysis

Analyze the antiviral effect of the filter material based on the observation results and assess its ability to inactivate the H1N1 virus.

#### Result Discussion

Discuss the potential applications of the filter material’s antiviral effect and its significance in the field of air filtration.

**This experiment enables the evaluation of the antiviral activity of the new virus-killing composite air filter material in a real-world environment, providing a scientific basis for its further application and development.**

## TEST REPORT

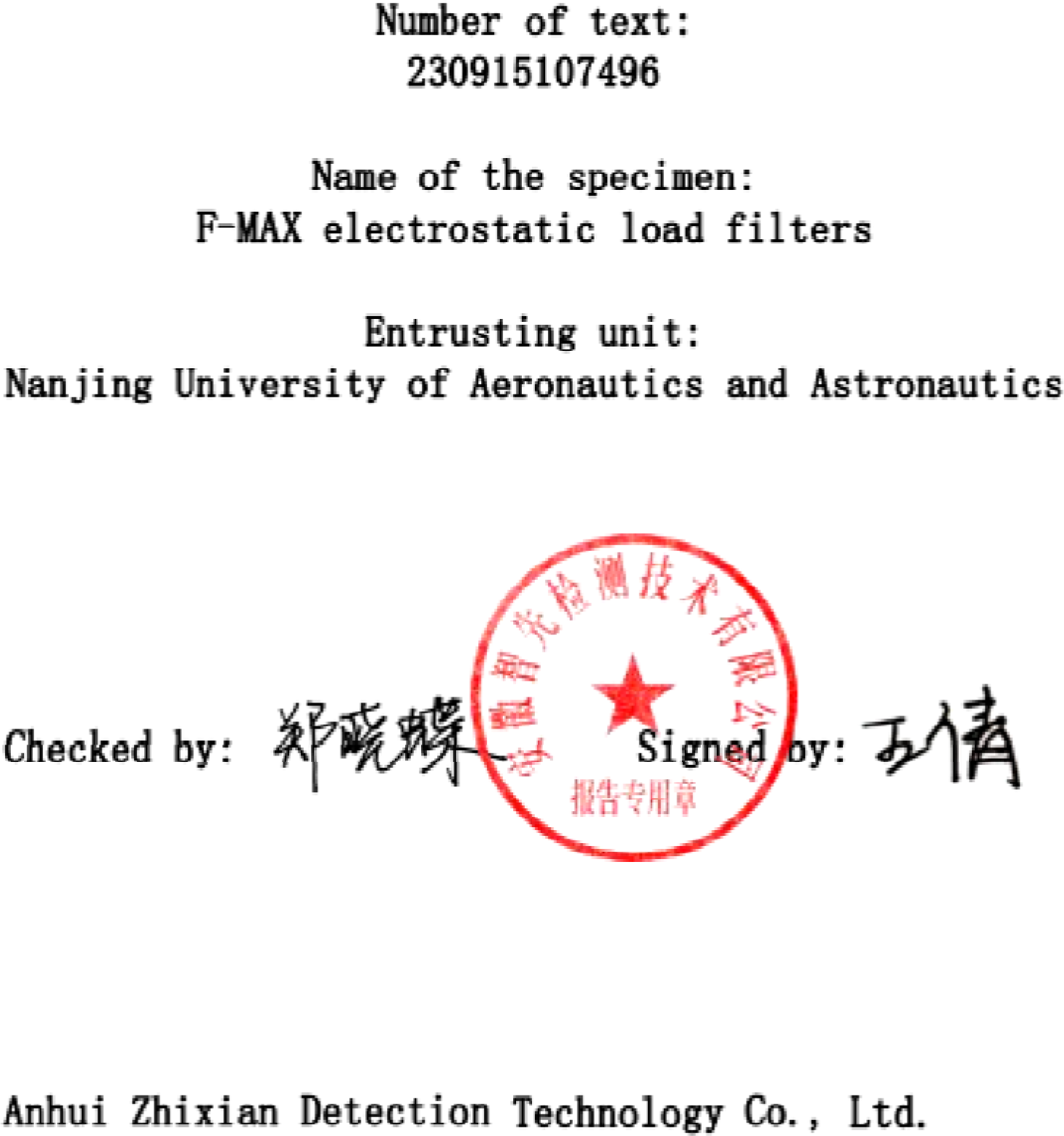

## ATTENTION

1. This report is invalid without the official seal of Anhui Zhixian Detection Technology Co., Lid.
2. This report is invalid without the signatures of the issuing and reviewing personnel.
3. Unauthorized modification, addition, or deletion of the report content renders the report invalid.
4. Without the consent of the company, this report may not be partially reproduced in any form; full reproduction is valid.
5. The testing results described in this report are only accountable to the commissioning entity for the provided samples.
6. If there are objections to the report results, they should be raised within 7 days from the date of receiving the report; any objections raised after this period will be considered as not valid.
7. The samples are generally retained for a period of 10 days after the completion of testing.
8. Our organization maintains confidentiality obligations for technical documents, conftact files, report texts, and other business secrets of the commissioning entity.
9. Reports without the CMA qualification accreditation mark are intended for research, teaching, and internal quality control purposes only and are not considered as data for social justice.
10. The content of the report in both Chinese and English is based on the Chinese version.

Company Information:

Company Name: Anhui Zhixian Detection Technology Co., Ltd.283 Fanhua Avenue, Shushan District, Hefei City, Anhui Province, China.

---

|  |  |  |  |
| --- | --- | --- | --- |
| Sample name | F-MAX electrostatic load filters | No. Of sample | / |
| Entrusting unit | Nanjing University of Aeronautics and Astronautics |  |  |
| Type of test | commissioned testing | Number of sample | 1 |
| Type of sample | / | Status of sample | Good |
| Testing period | 2023.09.16~2023.10.13 |  |  |
| Data of report | 2023.11.06 |  |  |
| Test | Virus test |  |  |
| Reference | ISO 18184:2019 |  |  |
| Experiment equipment | / |  |  |
| Test result | This report only provides actual measured values; detailed test results are available on the following page. |  |  |
| Other information | / |  |  |

| Virus | Host | Group |  | Viral infection titer log value (lgTCID<sub>50</sub>/mL) |  | Average viral infection titer (TCID<sub>50</sub>/mL) | Anti-irus reactivity Mv | Rate of virus reduction (%) |
| --- | --- | --- | --- | --- | --- | --- | --- | --- |
|  |  |  |  | Result | Average |  |  |  |
| H1N1 | MDCK Cell | 0h | Ref1 | 5.20 | 5.18 | 1.5 × 10<sup>5</sup> | >4.68 | >99.99 |
|  |  |  | Ref2 | 5.00 |  |  |  |  |
|  |  |  | Ref3 | 5.33 |  |  |  |  |
|  |  | 24h | Ref1 | 4.33 | 4.40 | 2.5 × 10<sup>4</sup> |  |  |
|  |  |  | Ref2 | 4.43 |  |  |  |  |
|  |  |  | Ref3 | 4.43 |  |  |  |  |
|  |  |  | Exp1 | <0.50 | <0.50 | <3 |  |  |
|  |  |  | Exp1 | <0.50 |  |  |  |  |
|  |  |  | Exp1 | <0.50 |  |  |  |  |
|  |  | Ref of cell: Normally act |  |  |  |  |  |  |

\*\*\*REPORT END HERE\*\*\*

**Figure S1:**
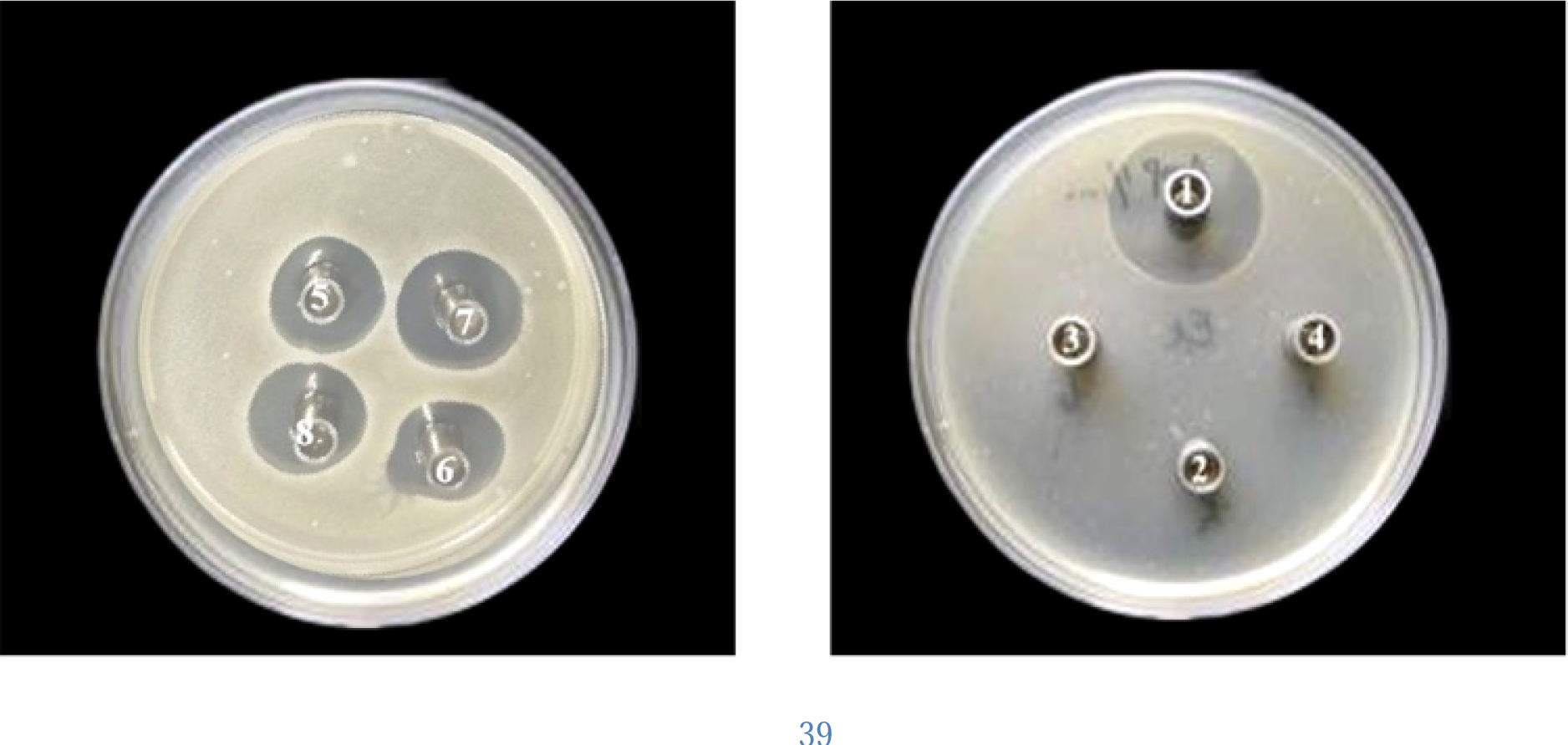
Antibacterial effect of F-MAX (antibacterial zone experiment).

**Figure S2:**
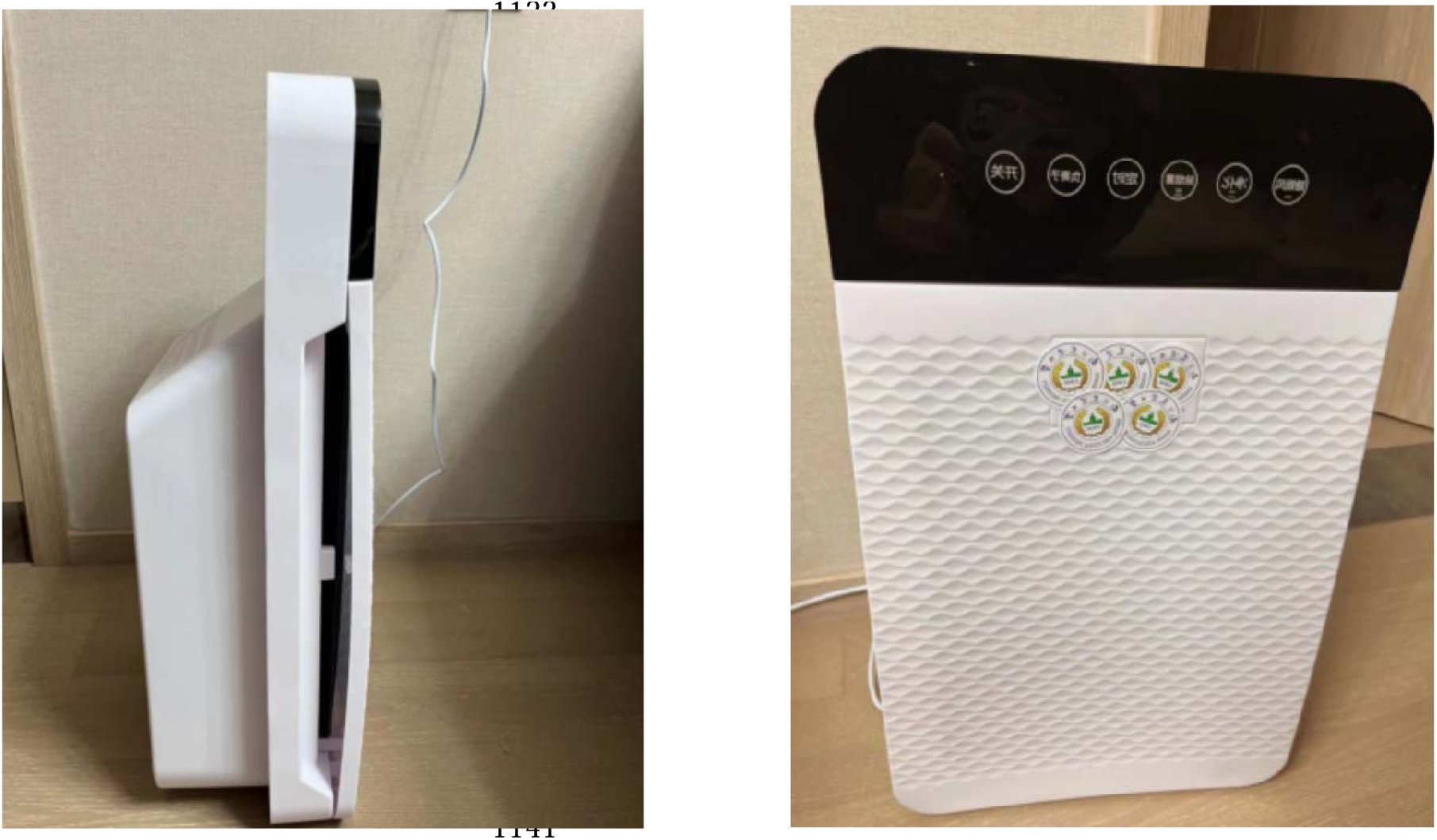
Photographs of T-robot.

**Figure S3:**
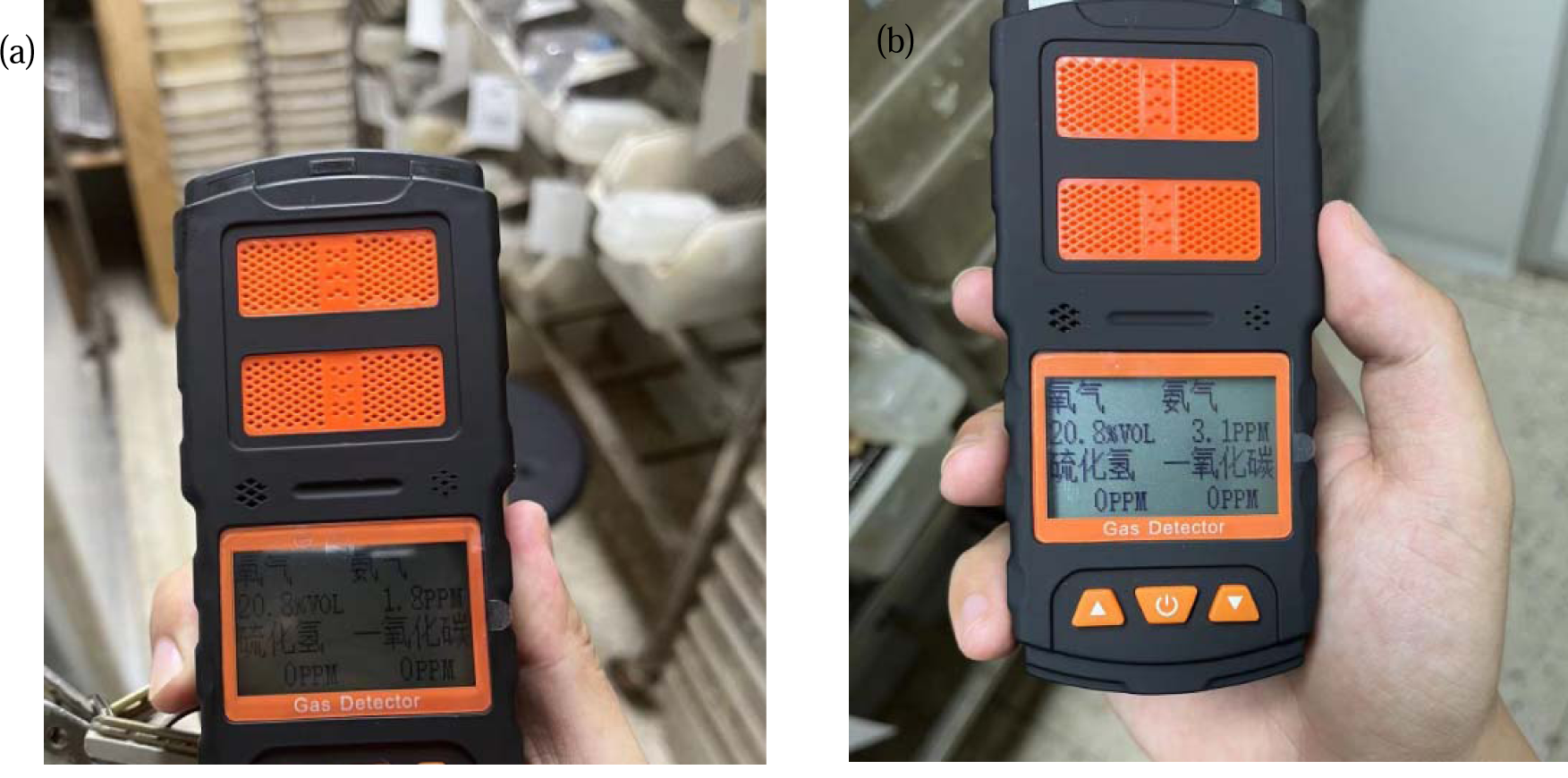
Ammonia filtration performance of the T-robot in a mouse room by showing Air condition of Mouse room before(a) and after (b) the use of T-ROBOT.

**Figure S4:**
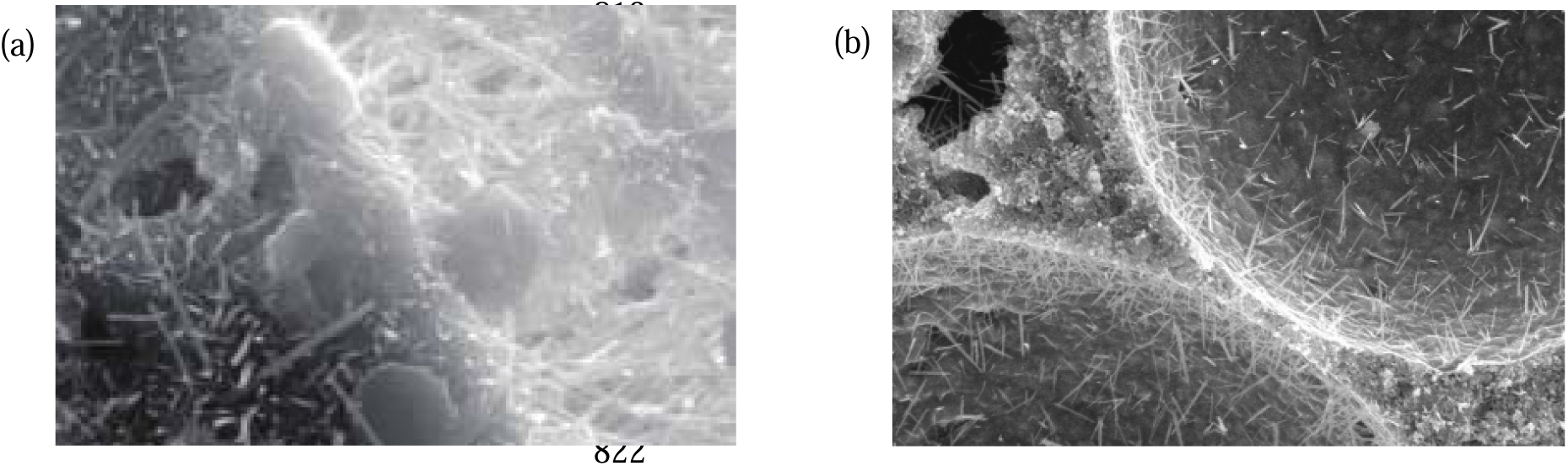
Comparison of SEM results of traditional(a) and newly developed(b) foaming BMSC paste.

**Figure S5:**
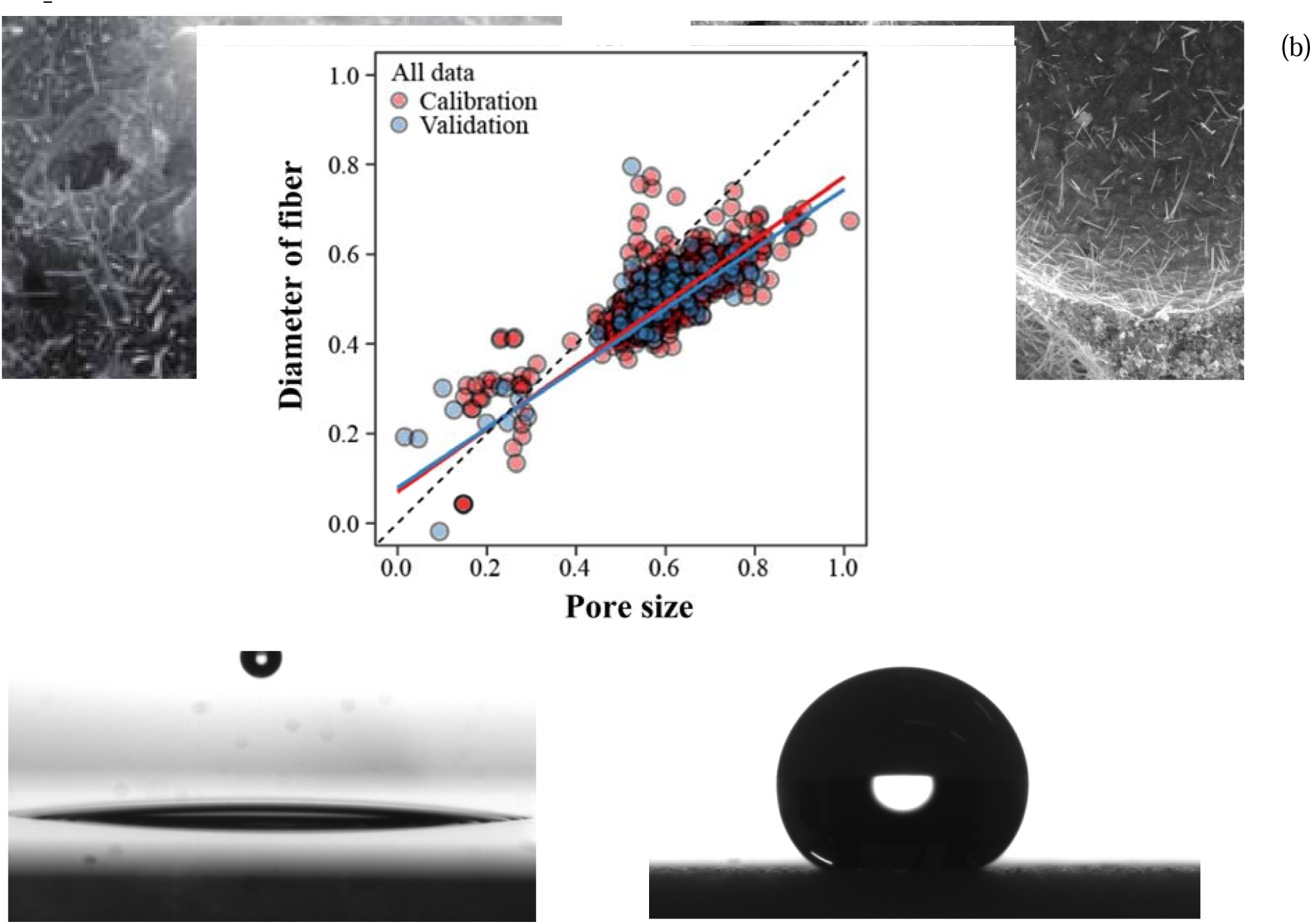
Experimental data using contact angle measurements before(a) and after(b) coating reveal significant hydrophobic enhancement. Initially hydrophilic.

**Figure S6:**
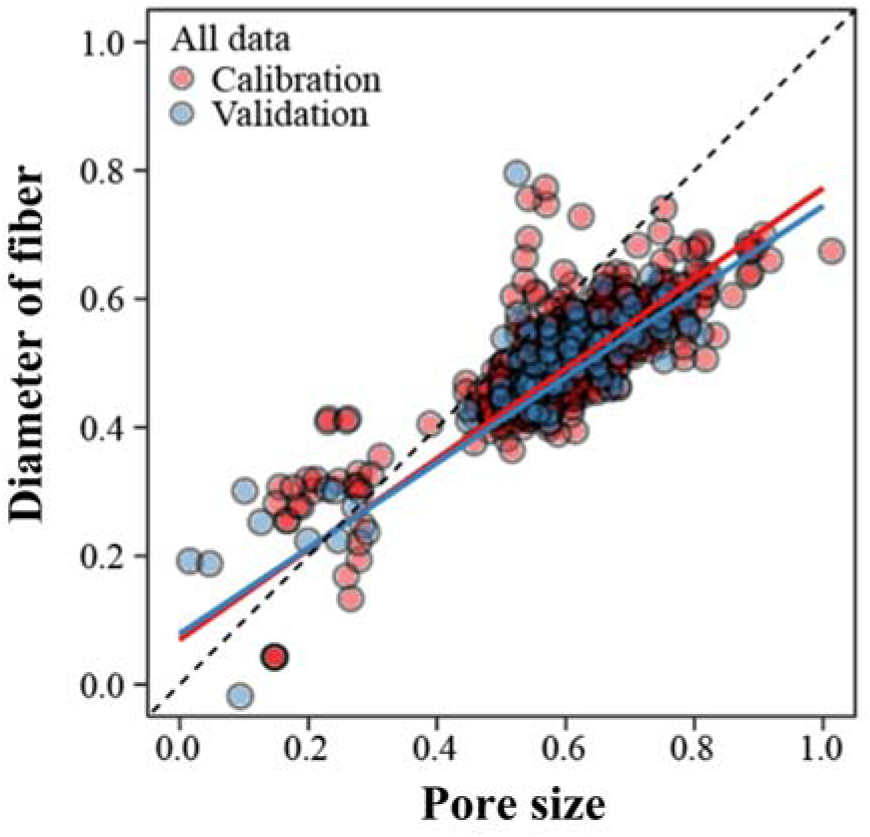
ML(Machine Learning) analysis for design of F-MAX electrostatically charged filter cartridge in the middle layer.

**Table S1:** Mix design of BMSC plate.

|  |  |  |
| --- | --- | --- |
| Mix design of BMSC plate (by mass) | MgO | 1 |
|  | MgSO<sub>4</sub> | 0.42 |
|  | Polycarboxylic acid additives | 0.016 |
|  | Water | 0.51 |
|  | Sand | 0.83 |
|  | Fly ash | 0.83 |

|  | Foaming agent | 0.01 |  |
| --- | --- | --- | --- |
|  | Specimen |  | Chinese<br>standard<br>JGJ/T17 |
| Strength,<br>MPa | Compressive strength, MPa | 7.8 | 3.50 |
|  | Splitting tensile strength, MPa | 2.5 | 0.56 |
|  | Tensile strength, MPa | 2.1 |  |
|  | Density, kg/m<sup>3</sup> | 156 |  |
|  | Elastic module, N/mm<sup>2</sup> | 2235 |  |

